# Communication from learned to innate olfactory processing centers is required for memory retrieval in *Drosophila*

**DOI:** 10.1101/167312

**Authors:** Michael-John Dolan, Ghislain Belliart-Guérin, Alexander Shakeel Bates, Yoshinori Aso, Shahar Frechter, Ruairí J.V. Roberts, Philipp Schlegel, Allan Wong, Adnan Hammad, Davi Bock, Gerald M. Rubin, Thomas Preat, Pierre-Yves Plaçais, Gregory S.X.E. Jefferis

**Affiliations:** Division of Neurobiology, MRC Laboratory of Molecular Biology, Cambridge CB2 0QH, UK; Janelia Research Campus, Howard Hughes Medical Institute, United States; Genes and Dynamics of Memory Systems, Brain Plasticity Unit, CNRS, ESPCI Paris, PSL Research University,75005 Paris, France; Department of Zoology, University of Cambridge, CB2 3EJ, UK

**Author notes:** Senior and Corresponding authors. Joint first author (Michael-John Dolan, Ghislain Belliart-Guérin).

## Abstract

Animals can show either learned or innate behavioral responses to a given stimulus. How these circuits interact to produce an appropriate behavioral response is unknown. In the *Drosophila* olfactory system, the lateral horn (LH) and the mushroom body (MB) are thought to mediate innate and learned olfactory behavior respectively, although the function of the LH has not been directly tested. Here we identify two LH cell-types (PD2a1/b1) that receive input from an MB output neuron required for recall of aversive olfactory memories. In contrast to the model above we find that PD2a1/b1 are required for aversive memory retrieval. PD2a1/b1 activity is modulated by training, indicating that memory information is passed to the innate olfactory processing centre. We map the connectivity of PD2a1/b1 to other olfactory neurons with connectomic data. This provides a circuit mechanism by which learned and unlearned olfactory information can interact to produce appropriate behavior.

## Introduction

The action of natural selection on evolutionary timescales can endow animal species with behavioural responses to stimuli of particular ethological relevance. In addition most animals also show adaptive responses based on learning during the lifetime of an individual. In some cases learning will potentiate or suppress an unlearned response. However, it remains unknown how memory recall interacts with innate sensory representations to produce the most appropriate behavior. This study explores this general issue using the *Drosophila* olfactory system. Olfaction is a shallow sense (in terms of layers of neural processing) with a privileged connection to memory systems in many species (Su et al., 2009). The genetic tractability of *Drosophila* and the numeric simplicity of its brain make it an ideal model to study how these processes interact at a neural circuit level (Masse et al., 2009), while the similarity of peripheral olfactory circuits across the animal kingdom makes it possible that neurobiological principles may also be shared deeper in the brain between insects and mammals (Su et al., 2009; Wilson, 2008).

In *Drosophila*, olfactory sensory neurons project to specific glomeruli in the antennal lobe (Masse et al., 2009; Su et al., 2009; Vosshall et al., 2000). Following local computations, excitatory uniglomerular projection neurons of the medial antennal lobe tract (PNs) make divergent connections to two higher processing regions, the lateral horn (LH) and the mushroom body (MB) (Keene and Waddell, 2007; Masse et al., 2009), in addition to other AL outputs (Strutz et al., 2014; Tanaka et al., 2012). The prevailing model of olfactory processing describes a clear functional division between these regions: the MB is required for learning, consolidation and retrieval of olfactory memories while the LH is thought to mediate innate behavior (Keene and Waddell, 2007; Masse et al., 2009). Many studies have confirmed the necessity of the MB for associative memory, where a reward or punishment (the unconditioned stimulus, US) is associated with one odor (the conditioned stimulus, CS+) but not with a second odor (CS-) (Keene and Waddell, 2007). The MB’s function in olfactory memory appears conserved in other insects such as ants (Vowles, 1964) and honeybees (Erber et al., 1980). The role of the LH in innate behavior has been inferred from experiments that silenced the MB and observed innate olfactory responses (Heimbeck et al., 2001; Parnas et al., 2013) (but see (Wang et al., 2003)). However, no studies to date have directly examined the behavioral functions of LH neurons in olfaction.

Brain mapping studies have shown that PNs from different glomeruli have stereotyped axonal projections in the LH (Datta et al., 2008; Jefferis et al., 2007; Kazama and Wilson, 2009; Marin et al., 2002; Wong et al., 2002) consistent with a role in innate olfactory behaviors. Anatomical and physiological analyses have shown a role for specific *Drosophila* LH neurons in processing pheromone cues likely relevant for sex-specific behaviors such as courtship and aggression (Jefferis et al., 2007; Kohl et al., 2013; Liang et al., 2013; Ruta et al., 2010). Recent results have shown that some LH neurons can also show stereotyped responses to general olfactory stimuli (Fişek and Wilson, 2014; Strutz et al., 2014) and are stereotypically connected to input PNs (Fişek and Wilson, 2014). In addition, new large scale data have confirmed response stereotypy and showed that different LHNs have wide variations in odor tuning and may encode odor categories (Fişek and Wilson, 2014; Kohl et al., 2013; Strutz et al., 2014) (SF and GSXEJ, personal communication).

In contrast to the LH, the neurons of the MB assembly are extremely well-characterised (Aso et al., 2014a). The dendrites of intrinsic MB neurons (Kenyon cells) are localised to a region called the calyx, where they sample incoming PN axons in an apparently random manner (Caron et al., 2013). Kenyon cells have parallel, axonal fibers which form five different lobes, with three distinct branching patterns that define as many Kenyon cell types (Aso et al., 2014a; Crittenden et al., 1998). A comprehensive anatomical analysis revealed that the lobes can be subdivided into 15 compartments, each innervated by specific dopaminergic input neurons (DANs) and MB output neurons (MBONs) (Aso et al., 2014a). These compartments are both anatomically and physiologically distinct (Cohn et al., 2015; Hige et al., 2015a), although each Kenyon cell axon forms synapses in all compartments of each lobe (Cohn et al., 2015).

Odors are sparsely represented in the Kenyon cell assembly so that in the lobes only a subset of axons will release neurotransmitter upon olfactory stimulation (Honegger et al., 2011; Perez-Orive et al., 2002; Szyszka et al., 2005). Electric shock, the US during aversive learning, activates a subset of DANs such that when US and CS+ are presented coincidentally, the subset of olfactory-driven Kenyon cells also receive dopaminergic input within specific compartments. This coincident input produces compartment-specific synaptic plasticity (Bouzaiane et al., 2015; Cohn et al., 2015; Hige et al., 2015a; Owald et al., 2015), changing the response of that compartment’s MBON to the CS+. MBONs function in valence behaviors, and a modified response to the trained odor may bias the fly’s behavior towards avoidance or attraction depending on the compartment (Aso et al., 2014b; Owald et al., 2015). One of these output neurons, MBON-α2sc (also known as MB-V2α), projects from the MB to several brain regions, including the LH (Séjourné et al., 2011). Previous work demonstrated MBON-α2sc is required for the retrieval of aversive olfactory memories across short, medium and long timescales (Hige et al., 2015a; Séjourné et al., 2011), although not necessary for the recall of appetitive memories (Felsenberg et al., 2017; Séjourné et al., 2011). Recordings from MBON-α2sc demonstrated it is broadly responsive to odor (Hige et al., 2015b) and depresses its response to the CS+ after training (Hige et al., 2015a; Séjourné et al., 2011). While stimulation of the entire V2 cluster (MBON-α2sc, MBON-α′3m and MBON-α′3ap) drives approach, activation of MBON-α2sc alone does not lead to any change in valence behavior (Aso et al., 2014b). Given the presumed role of the LH in innate olfaction, the function of this MB to LH projection is unclear. Is memory information transmitted to the LH and if so, is this communication required for retrieval of the aversive memory?

In this study we examine the behavioral function of this connection between the presumed innate and learned olfactory processing centers. We use a combination of computational anatomy and high-resolution microscopy to identify two LH output neuron cell-types (PD2a1 and PD2b1) which are postsynaptic to MBON-α2sc. We verify this connectivity with connectomic neuronal reconstruction. We then proceed to test the function of these cell-types in behavior. Contrary to the model of olfactory processing described above, PD2a1/b1 is necessary for memory retrieval. To confirm our observations we generated split-GAL4 lines (Luan et al., 2006; Pfeiffer et al., 2010) specifically targeting these neurons and used these reagents to confirm their necessity for memory recall. We performed calcium imaging of olfactory responses and similarly to MBON-α2sc, PD2a1/b1 also depress their responses to the CS+ after training. Using a whole-brain electron microscopy dataset (Zheng et al., 2017), we identify the PN input to PD2a1/b1 dendrites and show that these cells receive inputs from glomeruli associated with food and appetitive odors. We then show that PD2a1/b1 neurons potentially synapse back onto the MB network by interdigitating with select MBON axons and DAN dendrites implicated in approach behaviour and memory. Finally, we demonstrate that PD2a1/b1 forms axoaxonic synapses with MBON-α’2, an MBON necessary for the retrieval of appetitive memory (Aso et al., 2014b). This work provides a model for the interaction of innate and learned sensory information.

## Results

### Identifying neurons postsynaptic to MBON-α2sc in LH

In order to understand the role of information flow from the MB and LH, we first sought to identify the postsynaptic neurons in the LH that receive input from the MBON-α2sc. As no transsynaptic labelling system exists in flies, we developed a computational pipeline to find putative MBON-α2sc postsynaptic candidates in the LH. We used *in silico* overlap of GAL4 expression patterns to narrow down the field of candidate postsynaptic cell-types. Using image registration (Jefferis et al., 2007) of MBON-α2sc expressing a presynaptically localised marker (Christiansen et al., 2011), we created a mask of the MBON axonal terminals in the LH within template brain space. We then examined pixel overlap of the mask with registered images (Manton et al., 2014) of published GAL4 lines (Jenett et al., 2012). We ranked the lines by an “overlap score”, for which each brain was compared to GFP signal within the MB peduncle to exclude lines with MB expression. This allowed us to identify lines with no Kenyon cell expression, which could complicate behavioral analysis. The distribution of scores for approximately 3,500 GAL4 lines shows that the majority of lines lie close to zero or have negative scores (Figure 1A). Negative scores are derived from lines with little or no LH overlap and strong peduncle expression. We chose to focus on the top ∼100 lines, above the 0.97 quantile. After excluding lines driving expression in MBON-α2sc, examination of these top hits identified 5 cell-types putatively postsynaptic to the MBON-α2sc in the dorsal LH. As many lines were excluded during manual annotation due to broad expression, there are certainly other LH neurons we did not analyze in this study.

**Figure 1:**
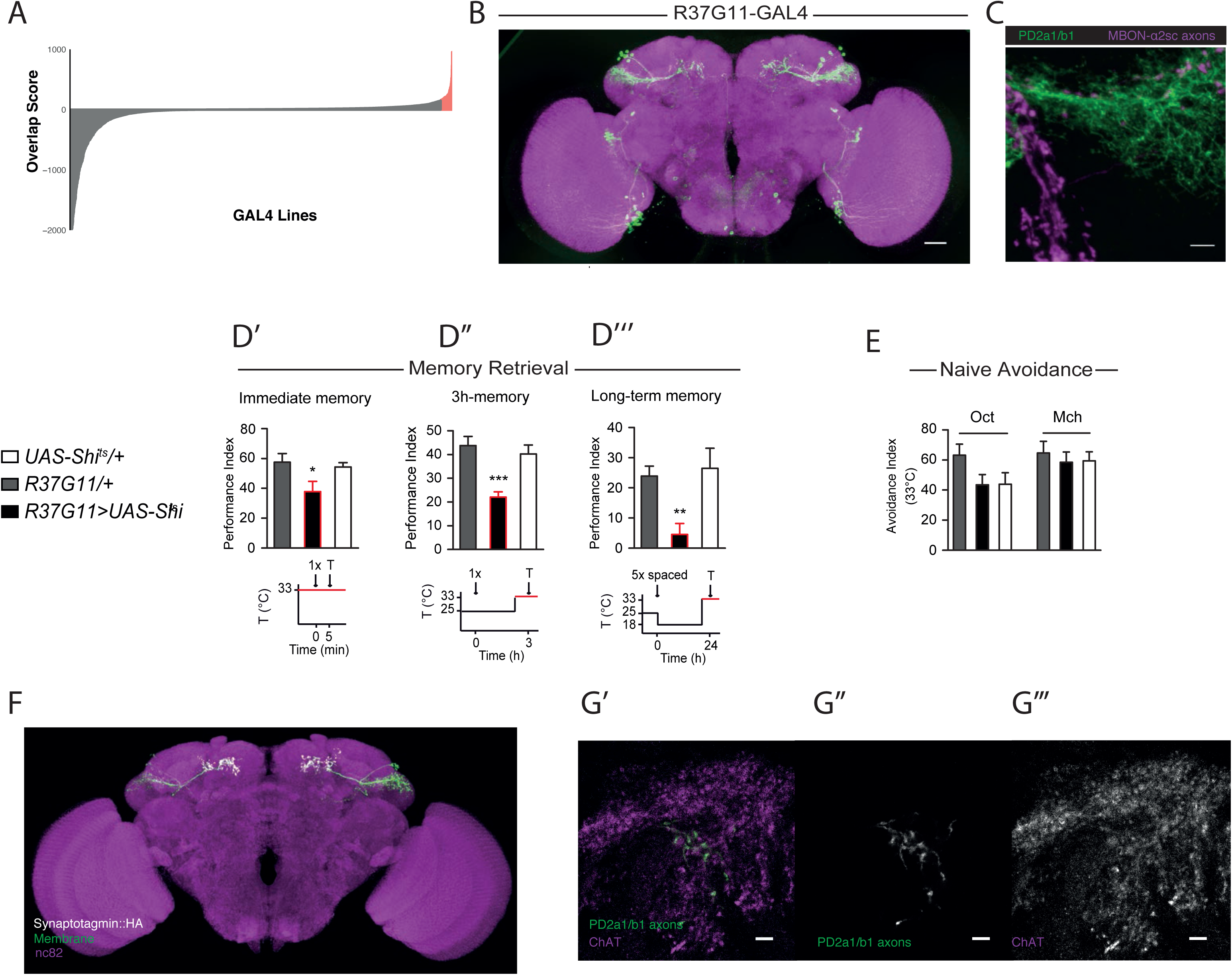
PD2a1/b1 is postsynaptic to MBON-α2sc and necessary for memory retrieval. (A) Summary distribution of overlap scores for the MBON-α2sc axon mask in the LH against all the GAL4 lines in the Janelia GMR collection. Lines that scored above the .97 quantile are labelled in red. Note that the Y axis is clipped below −2000 for clarity. (B) The sparsest GAL4 line labelling cell-type PD2a1/b1, R37G11-GAL4. Image stack is from the FlyLight website (www.janelia.org/gal4-gen1). The Scale bar is 30μm. (C) Z-projection of double labelling. Note that this LexA line contains both MBON-α2sc (dorsal) and MBON-α′3ap (ventral) axons are labelled in magenta while PD2a1/b1 is labelled with membrane-bound GFP (in green). The Scale bar is 5μm. Image is representative sample from n=4. (D) Flies expressing Shi_ts_ driven by R37G11-GAL4, and their genotypic controls, were trained and tested according to the illustrated protocols (periods at the restrictive temperature are indicated in red). Silencing PD2a1/b1 neurons impaired immediate memory after single-cycle (1x) training (D’: n = 12–13, F_(2,36)_ = 3.79, p = 0.033), 3h-memory after single-cycle training (D’’: n = 9, F_(2,26)_ = 12.07, p = 0.0002), and long-term memory after spaced training (D’’’: n = 9, F_(2,26)_ = 6.28, p = 0.0064). (E) Flies expressing Shi_ts_ driven by 37G11-GAL4 driver showed normal olfactory avoidance to octanol (Oct) and methyl-cyclohexanol (Mch) compared to their genotypic controls at the restrictive temperature (Oct: n = 14, F_(2,41)_ = 2.41, p = 0.10; Mch: n = 14, F_(2,41)_ = 0.23, p = 0.79). Data are presented as mean±SEM. (F) Confocal z-projection of PD2a1/b1 driving both membrane-bound GFP (green) and Synaptotagmin-HA (grey). PD2a1/b1 has been manually segmented in both channels. The image is registered with the neuropil marker nc82 (magenta) to the JFRC2013 template brain. Image is representative of n=5. (G’-G’’’) Representative single confocal slices of ChAT immunohistochemistry demonstrating that PD2a1/b1 neurons are cholinergic. Image is representative slice of n=4 stacks. The Scale bar is 5μm.

We next generated a LexA line to orthogonally control MBON-α2sc (Figure S1A), and conducted double-labelling of MBON presynapses and various LH cell-types identified in the computational screen to narrow down the number of identified cell-types. Two of the five cell-types had potential synaptic sites identified using double-labelling and high-resolution confocal microscopy: LH output neuron cell-types posterior dorsal 2a1/b1 (PD2a1/b1) (Figure 1B-1C, see below for single neuron data) and anterior ventral 6a1 (AV6a1) (Figure S2A, S2C). These names are defined by a hierarchical nomenclature for neurons with LH dendrite, which will be presented in a forthcoming manuscript (SF and GSXEJ, in preparation). This nomenclature considers cell body/primary neurite tract, primary dendrite and arborisation zones.

We also repeated this analysis for a mask of MBON axonal processes in the superior intermediate protocerebrum (SIP, data not shown) which yielded two clear, candidate postsynaptic cell-types, PD2a1 and PD2b1 (Figure S2B). This was further supported by double labelling (Figure S2D).

### PD2a1/b1 is necessary for memory retrieval

We identified the sparsest GAL4 lines for each of the three cell-types identified and screened for memory retrieval defects when the neurons were silenced in an aversive olfactory associative conditioning paradigm. LH cell-types expressed the temperature-sensitive neuronal silencer shibire^ts1^ (Kitamoto, 2001) which inhibits neuronal signalling at high temperatures (33°C, the restrictive temperature). By raising the temperature (from permissive 25°C to restrictive 33°C) during a memory test 3h after an aversive olfactory conditioning we could silence these neurons to probe their role specifically in memory recall (Séjourné et al., 2011).

Silencing the AV6a1 and SIP cell-type GAL4 lines had no detectable effect on memory performance (Figure S2G-H). However, silencing PD2a1/b1 neurons with R37G11-GAL4 impaired memory retrieval relative to genotype (Figure 1D’’) and permissive temperature (Figure S3B) controls in a 3h memory test assay. We extended these analyses of PD2a1/b1 to include immediate and long term memory which also require MBON-α2sc (Bouzaiane et al., 2015; Séjourné et al., 2011). Silencing PD2a1/b1 neurons attenuated memory retrieval for both phases of memory (Figure 1D’ and 1D’’’) compared to genotype and permissive temperature controls (Figure S3A and S3C). Surprisingly, inhibition of PD2a1/b1 had no effect on naive olfactory avoidance to either of the two training odors at the same concentration used in our memory assay (Figure 1E), indicating that the observed phenotype was not a defect in innate olfactory processing, the presumed function of LH neurons. These results indicate that PD2a1/b1 signalling during the test phase is necessary for memory recall.

We confirmed that PD2a1/b1 is primarily a LH output cell-type by expressing HA-fused synaptotagmin (Syt::HA) to label presynapses (Robinson et al., 2002) (Figure 1F). We also observed some presynapses in the presumptive LH dendrites (Figure 1F), although this has been observed for many different types of LH output neurons (MJD and GSXEJ, unpublished). We next determined the neurotransmitter profile of these cells using immunohistochemistry. PD2a1/b1 was ChAT-positive (Figure 1G’-G’’’) but GABA- and dVGlut-negative (Figure S1B-C) indicating that neurons in these cell-types are cholinergic LH outputs.

### Generation and characterization of cell-type specific split-GAL4 lines

Although a relatively specific GAL4 line, R37G11-GAL4 contained several other cell-types that could confound our behavioral results. To confirm that PD2a1/b1 was indeed responsible for the memory retrieval deficit, we generated split-GAL4 lines (Luan et al., 2006; Pfeiffer et al., 2010) that specifically labelled PD2a1/b1 neurons in the central brain (Figure 2A-B). Two of the lines identified, LH989 and LH991, used the R37G11-DBD hemidriver, the same enhancer that drove the PD2a1/b1 GAL4 line tested above. We focused on these two split-GAL4 lines because they were most likely to contain identical LH neurons implicated in GAL4 behavioral analysis. Both of these split-GAL4 lines also labelled neurons in the ventral nerve cord (VNC), however these VNC cell-types were different between lines (Figure 2A-B). We compared the number of PD2a1/b1 neurons labelled by each line. R37G11-GAL4 labelled 6.9±0.6 cells while LH989 and LH991 contained 5.25±0.5 and 5.67±0.8 neurons respectively.

**Figure 2:**
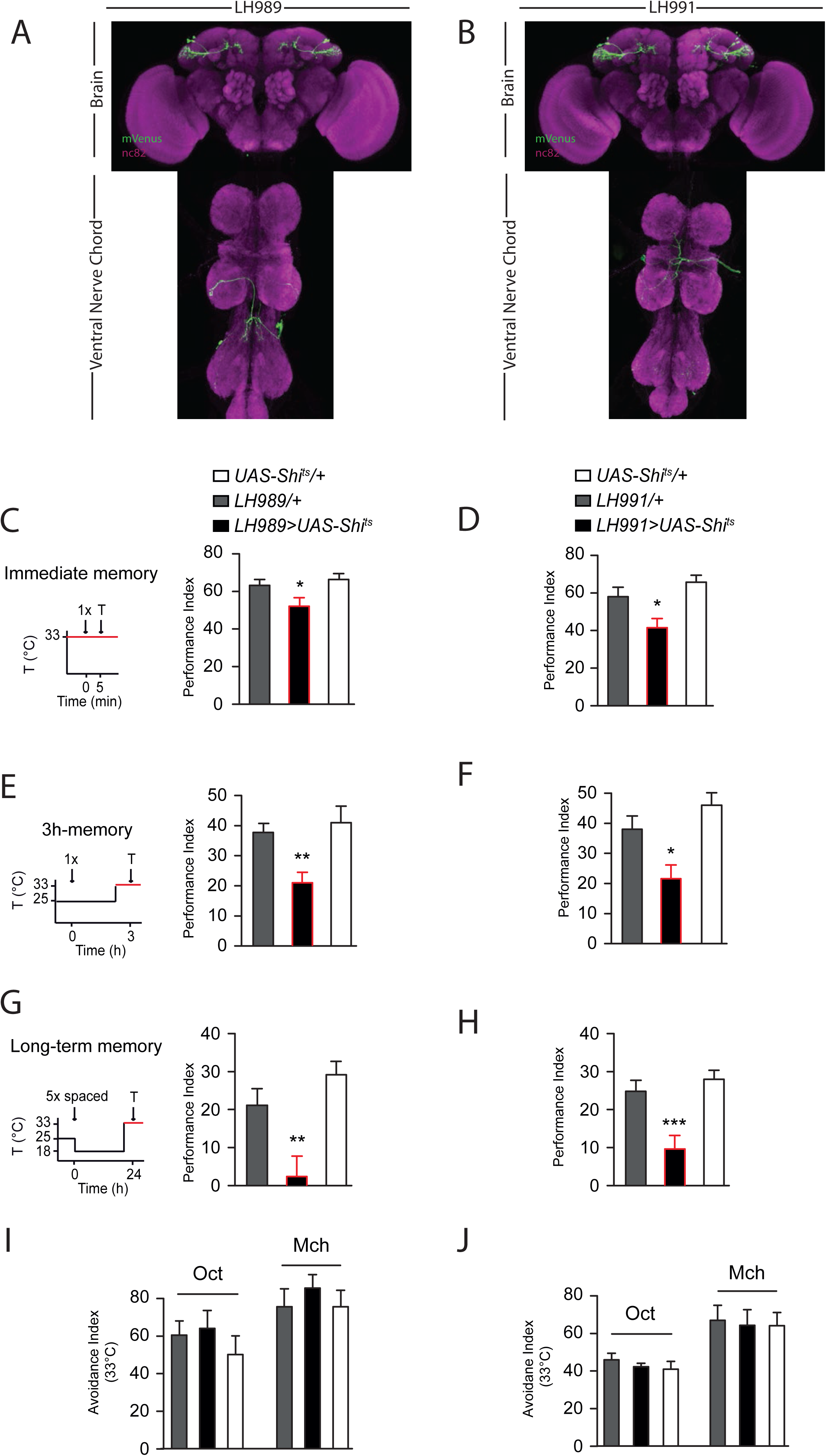
Specific control with the split-GAL4 system confirms PD2a1/b1’s role in memory retrieval but not innate behaviour. (A-B) Confocal z-projections of the two split-GAL4 lines generated to target PD2a1/b1 neurons, (A) LH989 and (B) LH991. Images contain both the brain and ventral nerve chord. Green is the membrane stain (mVenus) while the neuropil immunolabelling is in magenta. Image is representative of n=2 for each genotype. Flies expressing Shi_ts_ through the split-GAL4 lines LH989 or LH991 were trained and tested according to the illustrated protocols along with their respective genotypic controls (periods at the restrictive temperature are indicated in red). (C-D) Silencing PD2a1/b1 neurons using LH989 (C: n = 14–15, F_(2,42)_ = 4.13, p = 0.02) or LH991 (D: n = 18, F_(2,53)_ = 7.27, p = 0.0017) impaired immediate memory after 1x training. (E)-(F) Silencing PD2a1/b1 neurons during the retrieval phase 3h after 1x training using LH989 (E: n = 14, F_(2,42)_ = 6.73, p = 0.0031) or LH991 (F: n = 11–13, F_(2,35)_ = 8.23, p = 0.0013) caused a memory defect. (G-H) Silencing PD2a1/b1 neurons during the retrieval phase 24h after spaced training using LH989 (G: n = 7–9, F_(2,23)_ = 9.79, p = 0.0010) or LH991 (H: n = 19–23, F_(2,72)_ = 10.83, p < 0.0001) abolished performance. (I-J) Silencing PD2a1/b1 neurons using LH989 (I: Oct: n = 8–12, F_(2,29)_ = 0.63, p = 0.54; Mch: n = 10, F_(2,29)_ = 0.44, p = 0.65) or LH991 (J: Oct: n = 7–8, F_(2,22)_ = 0.25, p = 0.78; Mch: n = 7, F_(2,20)_ = 0.068, p = 0.93) had no effect on naive avoidance of Oct or Mch. Data are presented as mean±SEM.

To confirm that PD2a1/b1 is involved in the retrieval of several memory phases, immediately after single-cycle training, on the middle-term time scale (∼3h), and 24h after spaced training, we repeated our behavioral experiments with these sparse split-GAL4 lines. When flies were tested at the restrictive temperature to silence PD2a1/b1, memory performance was impaired in each of these three conditions compared to genotype controls (Figure 2C-H). This ranged from mild attenuation immediately after training (Fig. 2C,D) to full impairment for LTM retrieval (Fig. 2G,H), similar to phenotypes observed by silencing MBON-α2sc (Bouzaiane et al., 2015; Séjourné et al., 2011). This defect was due to neuronal silencing, as genotypically identical flies tested at the permissive temperature had no memory recall deficits at any phase (Figure S4). Finally we confirmed with these sparse split-GAL4 lines that silencing these PD2a1/b1 neurons had no effect on innate olfactory avoidance for the two training odors in our behavioral assay (Figure 2I-J), indicating that the observed defect was specific to memory recall. Output from cell-type PD2a1/b1 is therefore necessary for the retrieval of aversive olfactory memory, with the same characteristics as MBON-α2sc.

To understand the anatomy of the constituent cells within the PD2a1/b1 cell-type, we labelled single neurons in R37G11-GAL4 and the two split-GAL4 lines using the MultiColor FlpOut (MCFO) (Nern et al., 2015) technique (Figure S5A-C), isolating 22 single neurons from the PD2a1/b1 cell-type. 3/22 labelled neurons also projected to the MB calyx (and indeed this was visible in projection patterns of R37G11-GAL4, LH989 and LH991), while all other neurons appeared indistinguishable (Figure S5B-D). Therefore these lines label two distinct cell-types, PD2a1 (without calyx projections) and PD2b1 (with calyx projections). The calyx is the site of PN input to the MB, upstream from the site of associative olfactory memory, arguing against a role for this connection in our memory retrieval phenotype. As we could not get specific genetic control of these groups we refer to these cells as PD2a1/b1. PD2a1/b1 neurons appear morphologically similar to some members of a broader class of Type I LH neurons identified in a previous study (Fişek and Wilson, 2014).

### MBON-α2sc drives activity in PD2a1/b1

Our double labelling experiments suggested that MBON-α2sc is presynaptic to PD2a1/b1, but light microscopy does not have the resolution to confirm synaptic connectivity. We first used GRASP (Gordon and Scott, 2009) as an independent measure of the adjacency of PD2a1/b1 dendrites and MBON axons. The experimental genotype displayed clear GFP reconstitution in the dorsal LH (Figure 3A), indicating that the processes are proximal enough to form synapses, while no signal was detected in either of the control genotype brains (Figure 3B-C).

**Figure 3:**
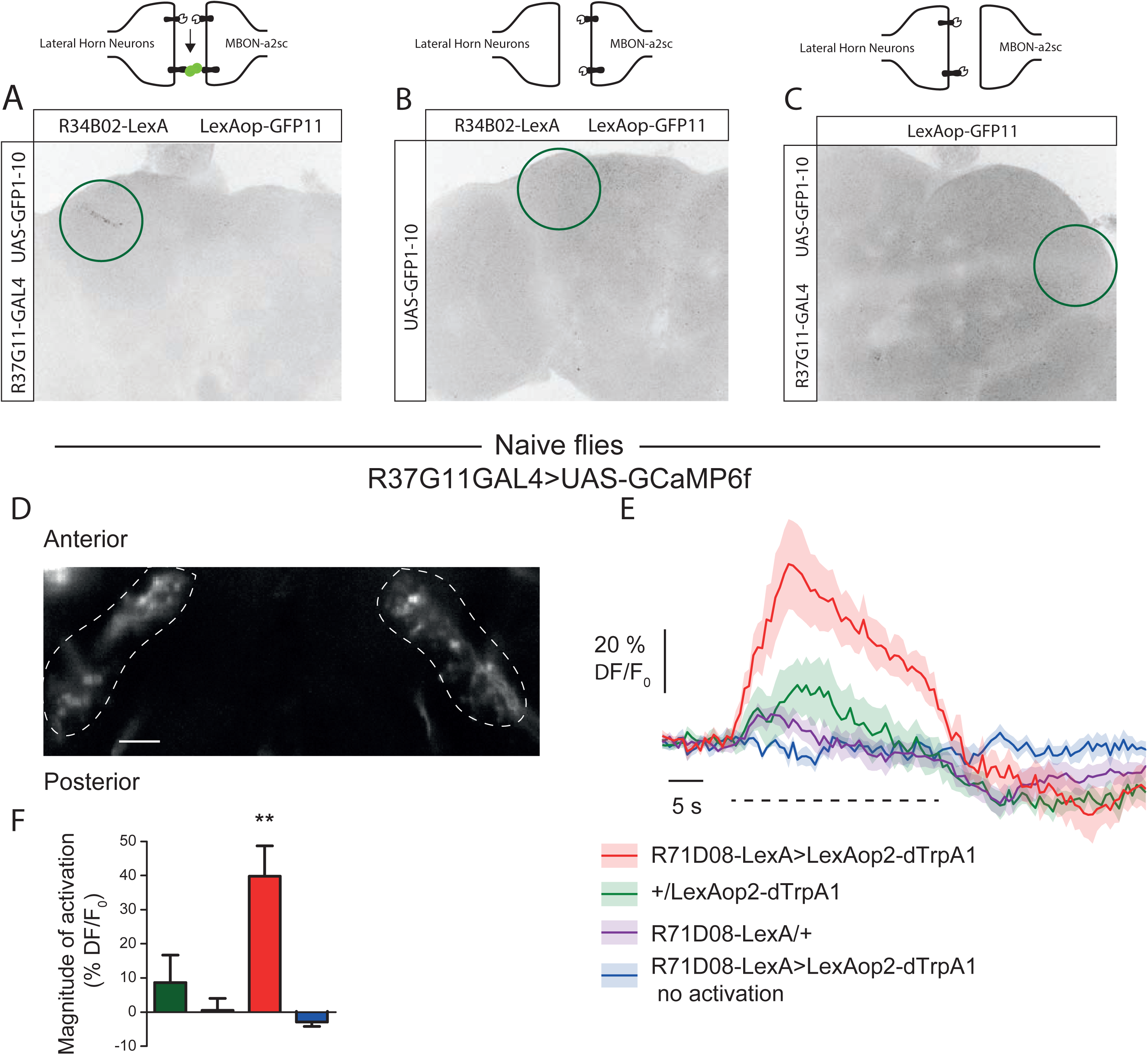
MBON-α2sc is functionally connected to PD2a1/b1. (A-C) GFP reconstitution across synaptic partners (GRASP) signal in the dorsal LH (indicated by green circle) for the experimental genotype (A), and two controls. (B-C) Genotypes are indicated and missing components are represented in the schematics above each figure. Images are representative of n=3 for controls and experimental genotypes. (D) GCaMP6f was expressed in PD2a1/b1 neurons with the R37G11-GAL4 driver (scale bar: 10μm). Fluorescence of the sensor was recorded in vivo from the axonal compartment of PD2a1/b1 neurons while the temperature of the imaging chamber was shifted from 20°C to 31°C (dashed line on F, except for the blue trace). The heat-sensitive channel dTrpA1 was expressed in V2 MBONs with 71D08-LexA. (E) The calcium increase of PD2a1/b1 neurons due to thermal activation of V2 MBONs (red trace) was stronger than that due to temperature shift only in the genotypic controls (green and purple trace). (F) Quantification of calcium increase from the traces shown in F (n = 10 flies per condition except 71D08-LexA/+ (n = 8), F_(3,37)_ = 9.09, p = 0.0001). Data are presented as mean±SEM.

As MBON-α2sc is cholinergic (Aso et al., 2014a; Séjourné et al., 2011), we would expect that stimulation of this neuron would drive activity in PD2a1/b1 if these neurons are connected. Initial experiments with optogenetic tools failed, as targeted expression of CsChrimson (Klapoetke et al., 2014) caused either fatality or miswiring in the two MBON-α2sc LexA lines available, mostly likely due to overexpression (data not shown). In an alternative approach we expressed the heat activated ion channel dTRPA1 (Hamada et al., 2008) to activate MBON-α2sc (Figure 3D-3F) while recording calcium transients in PD2a1/b1. We used R37G11-GAL4 to drive expression of GCaMP6f (Chen et al., 2013) and our R71D08-LexA line to drive dTRPA1 (Figure 3D). We imaged the axons of PD2a1/b1 *in-vivo* to determine if driving MBON-α2sc could induce calcium transients in PD2a1/b1. In a control experiment, we observed a small temperature-dependent increase in calcium in the absence of the LexAop2-dTRPA1 transgene indicating that temperature alone may also stimulate these neurons (Figure 3E-F), consistent with the projection of thermosensory neurons to the MB and LH (Frank et al., 2015). Similarly, we also observed a small calcium increase in flies carrying only the LexAop-dTRPA1 transgene (Figure 3E-F). However, increasing temperature in flies expressing dTRPA1 in MBON-α2sc yielded a larger increase in calcium indicating that these neurons are functionally connected (Figure 3E-F). We confirmed that dTRPA1 was expressing in MBON-α2sc by driving expression of a LexAop2-TdTomato reporter in the same landing site as the LexAop2-dTRPA1 transgene (Figure S6). This thermogenetic activation data, together with the double labelling and GRASP results suggest that MBON-α2sc connects to PD2a1/b1, LH cell-type that is necessary for memory retrieval.

### Synaptic resolution analysis of MBON-α2sc and PD2a1/b1 connectivity

A positive GRASP signal indicates that neuronal arbors of PD2a1/b1 dendrites and MBON-α2sc axons are in close physical proximity at some point in development, but does not conclusively demonstrate the existence of synaptic contacts. We therefore leveraged a new whole female brain serial section electron microscopy volume (Zheng et al., 2017) to examine anatomical connectivity with synaptic resolution. As there is only one MBON-α2sc per hemisphere (Aso et al., 2014a; Takemura et al., 2017), we first identified the MBON-α2sc on the ‘fly’s right’ in this brain volume, initially identifying it as a Kenyon cell postsynaptic target within the MB α2 compartment. We then used the NBLAST algorithm in conjunction with light-EM bridging registrations (Zheng et al., 2017) to match its backbone structure with previous light level image data (Aso et al., 2014a; Costa et al., 2016; Manton et al., 2014) (Figure 4A’-A’’). We repeated this analysis to identify the contralateral (left-side) MBON-α2sc as these neurons project bilaterally to both LHs. The reconstructed MBON-α2sc was split it into its axonic and dendritic compartments based on ‘synapse flow centrality’ (Schneider-Mizell et al., 2016).

**Figure 4:**
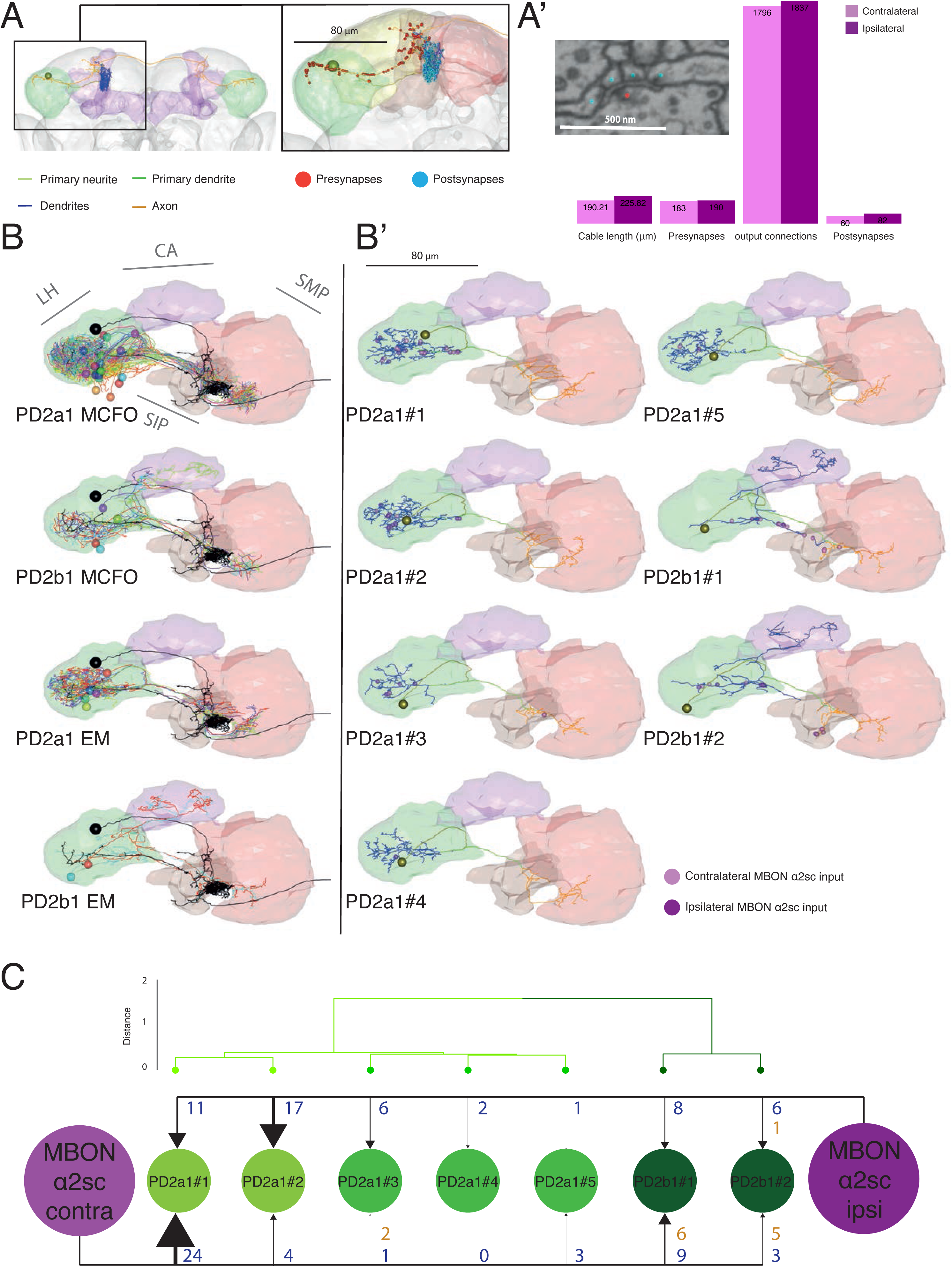
Electron Microscopy Reconstruction of PD2a1/b1. (A) Reconstruction of the right-side ‘ipsilateral’ MBON-α2sc in a whole brain EM volume. Cell body is represented as a sphere and the primary neurite (yellow-green), primary dendrite (green), dendrites (blue) and axon (orange) compartments are separately colored. Neuropils: LH in green, MB in purple. Inset, position of presynapses (red spheres) and postsynapses (cyan spheres) on right-side MBON-α2sc. Neuropils: SLP in yellow, SIP in orange, SMP in red. (A’) Comparison of different metrics for the reconstructions of the contralateral and ipsilateral MBON-α2sc, solely within the LH volume (green in A). Inset, example of a polyadic synapse with single T-bar (red dot) and multiple postsynapses (blue dots), referred to as ‘output connections’ in bar chart. Scale bar is 500nm. (B) Dorsal view of co-registered PD2a1 and PD2b1 MCFO data (top two panels respectively) and EM reconstructions (bottom two panels respectively). Cells are individually colored. Ipsilateral MBON-α2sc in black. (B’) Dorsal view of single PD2a1 and PD2b1 neurons reconstructed in the EM volume. Yellow-green sphere represents the soma while ipsilateral and contralateral MBON-α2sc synaptic connections are represented as dark and light purples respectively. Note that some spheres occlude further synapses at a similar location. Heavily innervated neuropils shown. (C) Schematic of synaptic connectivity from the two MBON-α2sc neurons onto each PD2a1/b1 cell, across all neuropils. Blue numbers indicate axo-dendritic connections and orange indicates axo-axonic. The PD2a1/b1 cells are clustered according to the NBLAST score of their axonal and dendritic compartments (see methods), identifying the two main groups, PD2a1 and PD2b1. Numbers beside each arrow indicate the number of outgoing connections made onto PD2a1/b1 neurons dendro-dendritically (blue) and axo-axonically (orange). Contra=contralateral, ipsi=ipsilateral, LH=lateral horn, CA=mushroom body calyx, SIP=superior intermediate protocerebrum, SLP=superior lateral protocerebrum, SMP=superior medial protocerebrum.

Next we reconstructed the axonal arbor in the right-side LH to completion for both MBON-α2sc neurons, marking presynapses and postsynapses for both, and annotating the connections each presynapse makes in the right LH (Figure 4A’’). However, the MBON-α2sc axon also targets other neuropils (Figure S8A). We identified 183 and 190 presynapses for the left and right-side MBON-α2sc respectively in the right-side LH (Figure 4A’’). We sampled 25% of these polyadic synapses’ connections (see Figure 4A’’ inset) and identified 70 substantial target skeletons (> 300 μm of neurite cable; data not shown). Using the same computational approaches, we determined that 2 of these target neurons had the distinctive morphology of the PD2a1/b1 cells. Using these PD2a1/b1 cells as a template, we located the PD2 primary neurite tract (SF and GSXEJ, in preparation) and reconstructed neurons in the PD2a1/b1 primary neuron tract (Figure S7A) to identify a total of five PD2a1 (PD2a1#1-5) and two PD2b1 (PD2b#1-2) cells (Figure 4B’-B’’, and see Methods). Comparison of MCFO alongside EM data confirmed the identity of PD2a1 and PD2b1 neurons (Figure 4B’ and Figure S7B). This was corroborated by a clustering analysis of NBLAST score that indicated no clear separation between EM, FlyCircuit (Chiang et al., 2011) and MCFO data (Figure S7C). Axons, dendrites, primary dendrites and primary neurite tracts were identified by splitting each neuron by synapse flow centrality (Schneider-Mizell et al., 2016). For both PD2b1 neurons the calycal projections were exclusively dendritic (Figure 4B’’, Figure S8D). We confirmed the existence of these two types of neurons by clustering a NBLAST score derived from dendrite and axonal compartments, which yielded two distinct groups for PD2a1 and PD2b1 (Figure 4C and SC’). NBLAST revealed that PD2a1 neurons could be further subdivided into two groups, one of which (comprising PD2a1#1 and PD2a1#2) receiving greater absolute MBON input per neuron (Figure 4C). Consistent with observations in the larva (Schneider-Mizell et al., 2016) and in the adult visual system (Scott et al., 2003), the vast majority of postsynapses were found on microtubule-free lower order branches of these adult, central brain neurons (Figure S8C and S10C). Summary data for pre- and postsynaptic sites, in addition to cable length for both MBON-α2sc and PD2a1/b1 can be found in supplementary information (Figure S8). We found that PD2a1/b1 synapses exhibited no electron-dense, only clear-core, vesicles, indicating that they do not release catecholamine or peptide neurotransmitters (data not shown).

All PD2a/b1 cells received input from either the ipsilateral, contralateral or both MBON-α2sc projections (Figure 4C). The majority of neurons received input from both MBON-α2sc neurons (Figure 4C), though three only very weakly. While we did not observe a privileged placement of MBON-α2sc postsynaptic placement on PD2a1.b1 dendrites (Figure S10A) we note that some sub-branches of certain PD2a1/b1 neuron’s dendritic trees receive a high proportion of MBON-α2sc input (Figure S10B). However, we can not rule out the possibility of additional indirect input from MBON-α2sc to PD2a1/b1. In sum, these observations confirm that PD2a1/b1 neurons are a direct synaptic partner of MBON-α2sc in the LH and further validate the directionality of the synaptic interaction is as expected from anatomy (Figure 1F), GRASP (Figure 3A-C) and physiology (Figure 3D-F).

### PD2a1/b1 neurons have decreased responses to the CS+

After training, MBON-α2sc depresses its response to the conditioned stimulus (Hige et al., 2015a; Séjourné et al., 2011). Next we wanted to determine if PD2a1/b1 neurons, being downstream of MBON-α2sc, also modulates its response to the CS+. We drove expression of the calcium indicator GCaMP3 (Tian et al., 2009) in PD2a1/b1 axons (Figure 5A). First, we observed in naïve flies that PD2a1/b1 neurons were responsive to 3-Octanol (Oct) and 4-Methylcyclohexanol (Mch), the two odorants that are alternatively used as CS+ in our behavioral experiments (Figure 1D–E and Figure 2). PD2a1/b1 neurons displayed significantly higher response for Oct than for Mch. Since MBON-α2sc responds with similar magnitude to these two odors (Hige et al., 2015b; Séjourné et al., 2011), this likely reflects input from antennal lobe PNs to the LH.

**Figure 5:**
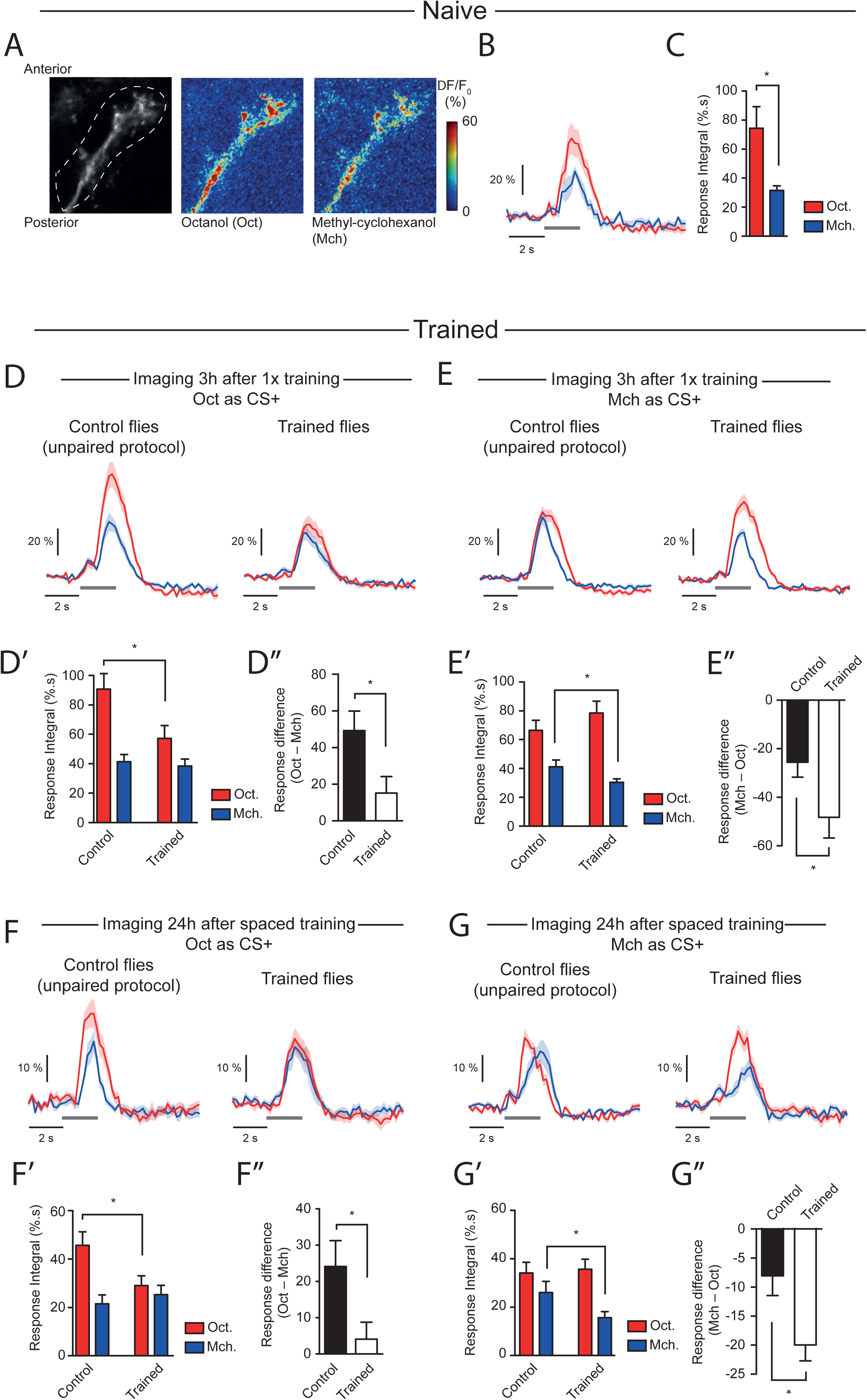
PD2a1/b1 decreases its response to the CS+ after training. (A) GCaMP3 was expressed in PD2a1/b1 with the R37G11-GAL4 driver. Olfactory responses to Oct and Mch were recorded in vivo from the axonal compartment of PD2a1/b1 neurons. (B-C) In naïve flies, the calcium increase in PD2a1/b1 neurons in response to Oct was larger than to Mch (average traces from n=6 flies; t-test, p = 0.015). (D) Odor responses were recorded 3h after 1x training using Oct as CS+ (n = 19 flies), or after the corresponding unpaired control protocol (n = 20 flies) (Fig. S9A).The integral of the odor responses (D’: t-test, p = 0.023) and the calculation of the difference between Oct and Mch responses (D’’: t-test, p = 0.024) revealed a decreased response to the CS+ after the associative protocol. (E) Odor responses were recorded 3h after 1x training using Mch as CS+ (n = 22 flies), or after the corresponding unpaired control protocol (n = 21 flies) (Fig. S7B). The integral of the odor responses (E’: p = 0.047) and the calculation of the difference between Mch and Oct responses (E’’: t-test, p = 0.041) revealed a decreased response to the CS+ after the associative protocol. (F) Odor responses were recorded 24h after 5x spaced training using Oct as CS+ (n = 9 flies), or after the corresponding unpaired control protocol (n = 11 flies) (Fig. S9A). The integral of the odor responses (F’: t-test, p = 0.036) and the calculation of the difference between Oct and Mch responses (F’’: t-test, p = 0.035) revealed a decreased response to the CS+ after the associative protocol. (G) Odor responses were recorded 24h after 5x spaced training using Mch as CS+ (n = 9 flies), or after the corresponding unpaired control protocol (n = 9 flies) (Fig. S9B). The integral of the odor responses (G’: t-test, p = 0.047) and the calculation of the difference between Mch and Oct responses (G’’: t-test, p = 0.010) revealed a decreased response to the CS+ after the associative protocol. Data are presented as mean±SEM. The grey bar indicates the period of olfactory stimulation. The delay between the switch of the valve and the onset of calcium response corresponds to the time for the odorant to flow from the valve output to the fly’s antennae. Although responses to Oct were systematically higher than to Mch (black bars in Figure 5D’’, E’’, F’’ and G’’), the magnitude of this difference varied depending on the sequence of odor presentation during conditioning, being lower when Mch was presented first. This occurred whether or not flies formed a memory, and may represent olfactory plasticity occurring in PNs or AL.

We next looked for training-induced alterations of odor responses, by comparing olfactory responses in PD2a1/b1 neurons following either associative training or a control unpaired protocol, which matched the odor sequence of the associative training but temporally separated electric shocks and odor delivery (See Figure S9 for schematic of full protocol). We performed these experiments either 3h after single-cycle training (Figure S9A-B), or 24h after spaced training (Figure S9C), and using either Oct or Mch as the CS+. Our measurements revealed that the pairing between the CS+ and electric shocks during single-cycle training resulted in a decreased CS+ response in PD2a1/b1 axons 3h later, either compared to unpaired controls (Figure 5D’ and E’) or relative to the CS-response in the same fly (Figure 5D’’and E’’). Similar results were observed 24h after spaced training, i.e. upon LTM formation (Figure 5F-G). This data is consistent with PD2a1/b1 neurons receiving memory-relevant information (the decreased CS+ response), resulting from depression at Kenyon cell to MBON-α2sc synapses.

### PD2a1/b1 also receives input from uniglomerular PNs encoding attractive odors

As noted above, PD2a1/b1 neurons are likely to receive input from PNs as well as MBON-α2sc. Indeed their dendrites are poised to integrate inputs from these two sources. A previous study identified morphologically similar neurons which are known to receive PN input (Fişek and Wilson, 2014). Since the uniglomerular, excitatory PN inputs to the LH and calyx have been mapped in the electron microscopy volume (Zheng et al., 2017), we sought to determine which of these PNs might provide input from the AL to PD2a1/b1 dendrites in the LH and calyx (Figure 6A’). In order to assess connectivity, we annotated presynapses in the LH for at least one PN from each glomerulus (n=73/112 PNs, 51 glomeruli; RJVR, PS, ASB, DB, GSXEJ, Scott Lauritzen, unpublished observations). For each PN observed to connect to a PD2a1/b1 neuron by at least one synaptic connection, we annotated presynapses for every PN of that glomerulus. We observed that most PD2a1/b1 neurons receive synaptic input from several glomeruli, chiefly DM1, DP1m, DM4, VA2, DP1l and VM3 (Figure 6A’), although differences were observed between the PD2a1/b1 cells. We combined our connectivity information with data on the number of sister PNs per glomerulus (Grabe et al., 2016) and calcium imaging of odor responses in PN dendrites (Badel et al., 2016). PN synapses were mostly located on lower order, microtubule-lacking branches (Figure S10A). We found that the strongest PN input onto PD2a1/b1 neurons was singlet PNs (ie. from glomeruli with only one PN, ∼21/53 right-side glomeruli in the present EM volume, RJVR, PS, ASB, DB, GSXEJ, Scott Lauritzen, unpublished observations) which responded sparsely to odorants (Figure S10B).

**Figure 6:**
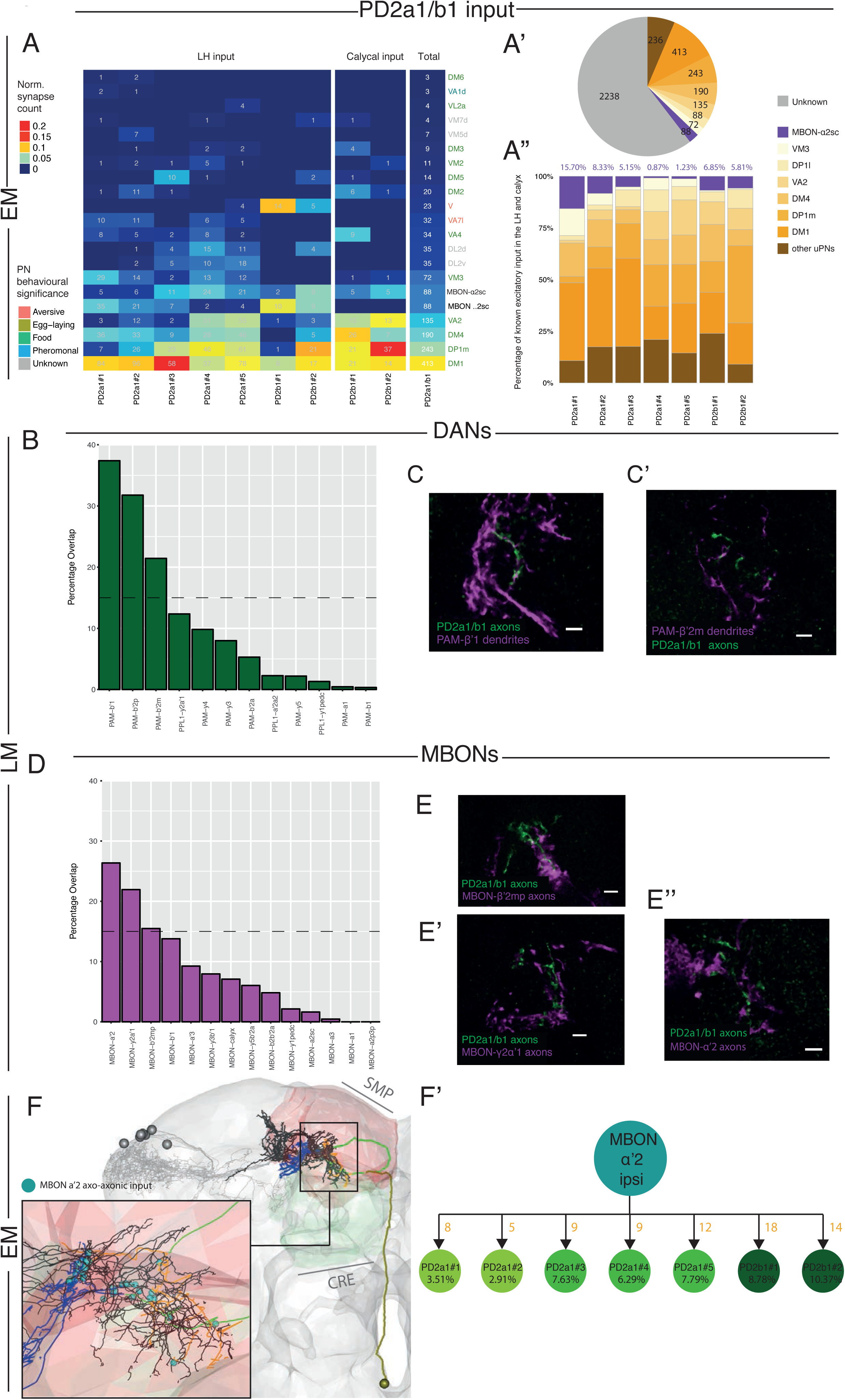
PD2a1/b1 receive input from appetitive PNs and converge with a subset of MBONs and DANs. (A’) Summary of antennal lobe glomeruli with uniglomerular, excitatory PN connectivity to PD2a1/b1 neurons. Uniglomerular PNs from connected glomeruli or MBON-α2sc ordered by connection strength (rows). Note that glomeruli whose uniglomerular PNs did not connect to PD2a1/b1 dendrites are not shown. Connectivity is divided by PD2a1/b1 dendrites in the LH, PD2b1 dendrites in the MB calyx and the sum total across all PD2a1/b1 dendrites in both compartments. A small number of dendritic postsynapses fell outside these neuropils and are not shown. Numbers represent the number of synapses while heatmap coloring represents the synapse count normalized by the total number of postsynapses the cell(s) have in the given compartment(s). (A’’) The different proportions of known excitatory synaptic input for each PD2a1/b1 dendrite. (A’’’) The number of synaptic inputs for all PD2a1/b1 dendrite traced in this study. Input is either undefined (gray), uniglomerular PN (oranges) or MBON-α2sc (purple). (B) Histogram of overlap between a mask of PD2a1/b1 axons and masks of the dendrites of most DANs (along the x-axis). (C’-C’’) High resolution double labelling of PD2a1/b1 axons (labelled with GFP, green) and DAN dendrites (labelled with RFP, magenta). (C’) PAM-β’1 dendrites (C’’) PAM-β’2m dendrites. (D) Histogram of overlap between a mask of PD2a1/b1 axons and masks of the dendrites of most MBONs (along the x-axis). (E’-E’’) High resolution double labelling of PD2a1/b1 axons (labelled with GFP, green) and MBONs (labelled with RFP, magenta). (E’) MBON-β’2mp axons, (E’’) MBON-y2α’1 and (E’’’) MBON-α′2 axons. (F’) Visualization of MBON-α’2 axons interdigitating with PD2a1/b1 axons (black) in the whole brain EM volume. The rest of the PD2a1/b1 neurons shown in gray. Inset, positions of axo-axonic connections onto PD2a1/b1 neurons shown as cyan spheres. (F’’) Summary of reconstructed ipsilateral MBON-α’2’s axo-axonic connectivity onto PD2a1/b1 cells. EM=Electron Microscopy, LH=Light Microscopy.

Interestingly, the majority of these glomeruli are responsive to appetitive and food odors (Badel et al., 2016; Knaden et al., 2012; Semmelhack and Wang, 2009) (Figure 6A), indicating that the PD2a1/b1 cell-type receives direct PN input mostly from appetitive olfactory channels. Input to both DM1 and VA2 glomeruli are required for approach behavior to vinegar (Semmelhack and Wang, 2009). Indeed the dorsal LH, where PD2a1/b1 dendrites are located, has previously been associated with coding of food odors (Jefferis et al., 2007).

We next examined direct synaptic connectivity in the calyx between Pd2b1 cells and PN boutons. Interestingly, these cells received input from the same four glomeruli as above (Figure 6A’). An exception was PD2b1#1, a cell which receives no DP1m or DM4 input in the lateral horn but receives synapses from these PNs in the calyx (Figure 6A’). PD2a1/b1 neurons were found to form presynapses in the LH, but not in the calyx (Figure S8).

The majority of input from PD2a1/b1 is undefined (Figure 6A’’-A’’’), and could be supplied by local neurons, other types of PNs, perhaps inhibitory PNs that ascend through the mediolateral antennal lobe tract and multi-glomerular PNs, or non-canonical LH input. This could also include indirect inputs to PD2a1/b1 dendrites from either PNs or MBON-α2sc. The PNs in aggregate make up the second-largest source of input while MBON-α2sc cells are responsible for 1.90% of total dendritic PD2a1/b1 postsynapses across these neurons (95 postsynapses, 88 in the LH and the rest elsewhere in the superior protocerebrum, Figure 4B’’) but 6.39% of known, direct excitatory input. Although relatively small, we note that MBON-α2sc has been shown to exhibit a sustained, high rate of firing after odor stimulation compared to PNs (Hige et al., 2015b; Wilson et al., 2004) and our thermogenetic experiments indicate that MBON-α2sc stimulation can drive activity in PD2a1/b1 (Figure 3D-F). In summary these data indicate that PD2a1/b1 receives both direct (from PNs) and indirect (from MBON-α2sc) olfactory input from the AL. This direct input consists mostly of neurons that respond to appetitive odors.

### PD2a1/b1 interdigitates with DAN dendrites and MBON axons in MB convergence zones

As a first insight into what role PD2a1/b1 may play in memory retrieval, we sought to identify potential downstream targets of this LH cell-type. Light and electron microscopy characterisation of PD2a1/b1 axons suggested it transmits information from the LH to the crepine (CRE), superior medial protocerebrum (SMP) and the superior intermediate protocerebrum (SIP) (Figure S8 and data not shown). The CRE, SMP and SIP have been identified as convergence zones that contain both the dendrites of DANs and the axons of MBONs (Aso et al., 2014a, 2014b; Owald et al., 2015). This raised the possibility that PD2a1/b1 may potentially interact with input and output neurons of the MB assembly.

We searched for potential sites of contact with MBONs and DANs by computational alignment of light microscopy data. We used segmented axon and dendrite masks in the same brain space (Jefferis et al., 2007; Rohlfing and Maurer, 2003), comparing PD2a1/b1 axons with published MB data (Aso et al., 2014a). We calculated the percentage overlap between the mask of PD2a1/b1 axons and either the axon or dendrite of each MB cell-type. This coarse analysis revealed several potential contacts between PD2a1/b1 and DAN dendrites (Figure 6B) and MBON axons (Figure 6D). We further investigated all neurons above a 15% threshold of PD2a1/b1 overlap using double labelling with R37G11-LexA, a PD2a1/b1 LexA line (Pfeiffer et al., 2010) (www.janelia.org/gal4-gen1).

We examined three DANs using double labelling. Both PAM-β′1 and PAM-β′2m interdigitated and exhibited potential synaptic contacts with PD2a1/b1 axons (Figure 5C’-C’’), suggesting potential synaptic contacts. PAM-β′2p had dendrites proximal to PD2a1/b1 axons but did not interdigitate (data not shown). PAM-β′1 mediates memory erasure (Shuai et al., 2015) and therefore a MBON-LH-PAM neuron circuit may play a role in forgetting or extinction. PAM-β′2m, together with PAM-β′2p can drive approach behavior when stimulated (Lewis et al., 2015).

Double labelling analysis of MBON axons and PD2a1/b1 axons revealed close co-projection for MBON-β’2mp, MBON-γ2α′1 and MBON-α′2 (Figure 5E’-E’’’), indicating common postsynaptic partners or possibly axo-axonic synapses. This data indicates that PD2a1/b1 projects to MB convergence zones, interdigitates with both DANs and MBONs, potentially feeding back onto the MB assembly (Aso et al., 2014a, 2014b). MBON-β’2mp receives input from the MB compartment that is innervated by PAM-β’2m and plays a role in appetitive and aversive memory retrieval (Owald et al., 2015). MBON-γ2α′1 drives approach when stimulated (Aso et al., 2014b), and has been implicated in appetitive memory retrieval; it is also necessary and sufficient for memory reconsolidation (Aso et al., 2014b; Felsenberg et al., 2017). Silencing of MBON-α′2 throughout training and testing abolishes appetitive memories (Aso et al., 2014b).

Convergence between PD2a1/b1 and MBON axons could imply either interdigitation or axo-axonic synapses. To test this and validate our light-level double labelling, we returned to electron microscopy and reconstructed the ipsilateral MBON-α′2, the MBON that gave the highest PD2a1/b1 axon overlap score for MBONs (Figure 6D). We discovered that MBON-α′2 makes axo-axonic connections onto PD2a1/b1 neurons (Figure 6F) indicating that PD2a1/b1 can be directly modulated by MBON-α′2. The close proximity between axonal arbors required to make multiple axo-axonic synapses further implies that PD2a1/b1 and MBON-α′2 may also share downstream targets. This illustrates that convergence with MBONs can imply synaptic connectivity and not just interdigitation. In aggregate these data suggests that PD2a1/b1 interacts or converges with MB neurons that drive approach behavior, memory retrieval and the reappraisal of memories (Figure 7A).

**Figure 7:**
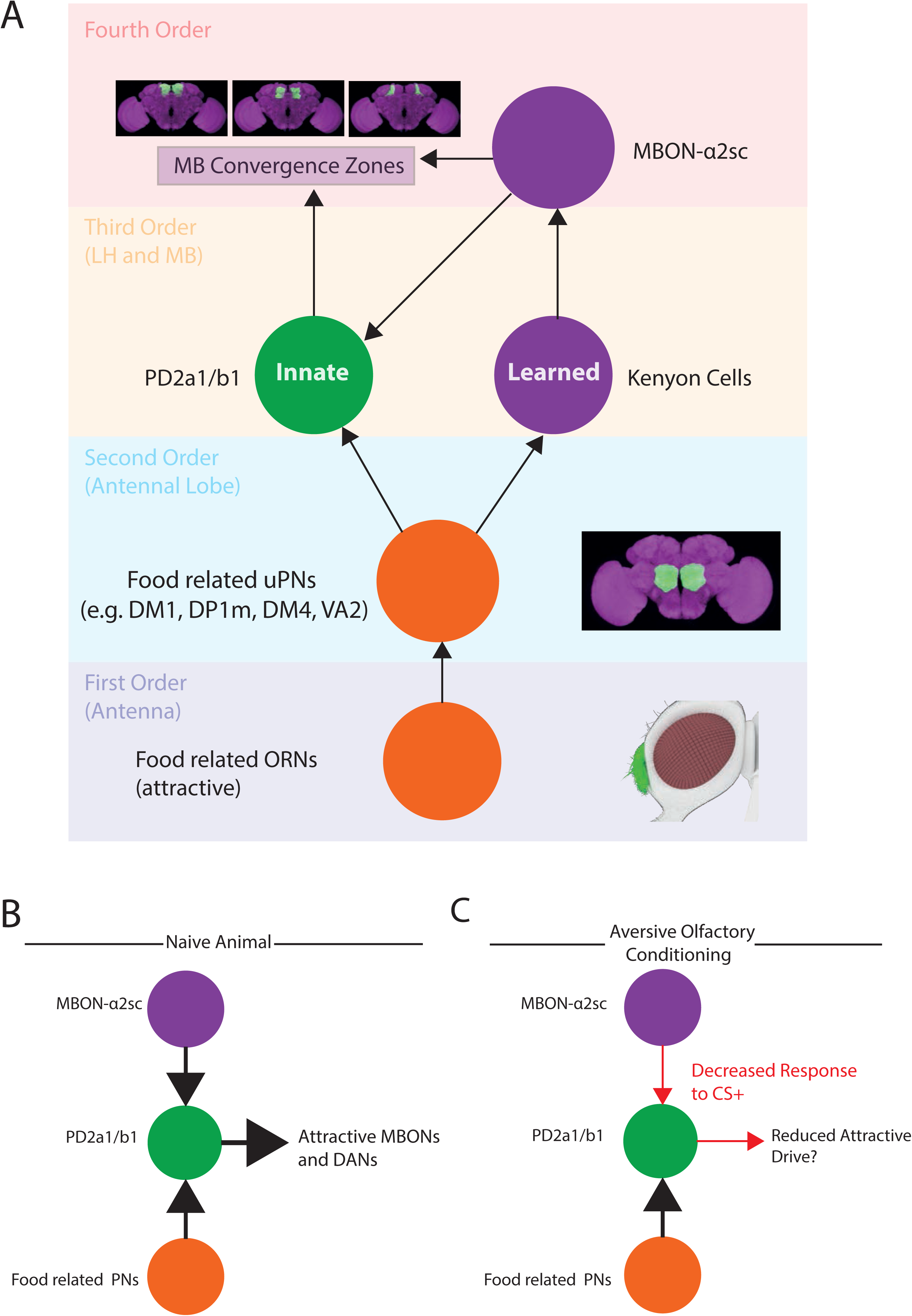
Summary and model of MB/LH interactions during memory retrieval. (A) Schematic illustrating the olfactory system of *Drosophila* and the interactions discovered in this study. Olfactory sensation occurs on the antenna and maxillary palps, the first order nodes of the olfactory system. ORNs project from these regions to the second order olfactory neuropil, the antennal lobe. Here we focus on the four uniglomerular PN types with the highest amount of synaptic input which encode food/appetitive odors. These PNs project to the third order nodes of the olfactory system, the MB and the LH. These PNs synapse onto both the kenyon cells of the MB and PD2a1/b1 dendrites within the LH (although they likely make contacts with other LH neurons too). MB output is read out by MBONs including MBON-α2sc, which projects to the LH and several regions in the fourth order relays of the olfactory system, and MBON-α’2. MBON-α2sc forms synapses onto PD2a1/b1 (and likely other LH neurons). PD2a1/b1 also projects to these fourth order neuropils where it converges with appeditive MBONs and DANs, thus interfacing with the input and output of the MB. The axons of MBON-α’2 form axo-axonic synapses onto the axons of PD2a1/b1. Note that the PN input to the dendrites of PD2a1/b1 in the MB calyx is not shown here. (B) Model for how PD2a1/b1 functions in naive and trained animals. In naive animals PD2a1/b1 receives input from both the MB (via MBON-α2sc) and food related PNs. PD2a1/b1 projects to DANs and MBONs implicated in appetitive and approach behaviors and this may drive or contribute to an odors net attractiveness. After conditioning the response of MBON-α2sc to the CS+ is reduced, resulting in a concomitant decreased response to the CS+ in PD2a1/b1. This reduces the input onto the appetitive DANs and MBONs which results in decreased attraction to the CS+.

## Discussion

Labelling and manipulating cells connected to genetically defined neuronal populations is challenging in *Drosophila*, in the absence of transsynaptic tracing tools. We developed a novel, simple and widely applicable approach to identify the putative postsynaptic partners of a given neuron (Figure 1A). By combining this analysis with double labelling, GRASP, thermogenetic mapping and neuronal reconstruction from electron microscopy we identify LH output neurons PD2a1/b1, neuronal cell-types directly postsynaptic to MBON-α2sc (Figure 7A).

Strikingly, using specific split-GAL4 control of PD2a1/b1 neurons in the brain, we found that PD2a1/b1 signalling is necessary for memory retrieval across all phases tested but dispensable for innate olfactory responses. It is possible that PD2a1/b1 neurons do participate in some innate olfactory behaviors, but they are not critical for the stimuli or context used in our memory tests. For the first time, to our knowledge, we have directly interrogated the role of LH neurons in olfactory behavior in adult *Drosophila,* discovering a LH cell-type that is necessary for memory retrieval, contrary to the assumption that the LH solely mediates innate behavior.

We next examined the responses of PD2a1/b1 to the training odors using *in-vivo* calcium imaging. Although odor tuning varies between individual flies, on average MBON-α2sc responds equally to both 4-Methylcyclohexanol and 3-Octanol (Hige et al., 2015b; Séjourné et al., 2011). PD2a1/b1 was also odor-responsive but unlike MBON-α2sc responded more strongly to Oct than Mch, as PD2a1/b1 integrates inputs from PNs and the MB. We observed that associative olfactory training induced a decreased response to the CS+ relative to the CS-in PD2a1/b1 neurons, in a manner similar to MBON-α2sc (Séjourné et al., 2011), indicating that memory information is sent from the MB to LH.

To determine the synaptic connectivity of PD2a1/b1 within the olfactory circuitry we leveraged a whole *Drosophila* brain electron microscopy volume (Zheng et al., 2017). We examined the connectivity of PD2a1/b1 dendrites with uniglomerular excitatory PN axons and found that PD2a1/b1 dendrites in the dorsal LH mostly receive input from PNs encoding appetitive/food odors (Hallem and Carlson, 2006; Knaden et al., 2012; Semmelhack and Wang, 2009), including uniglomerular PNs from the DM1 and VA2 glomeruli, which are necessary for attraction to vinegar (Semmelhack and Wang, 2009). Using a combination of light microscopy and electron microscopy reconstruction we demonstrate that PD2a1/b1 interdigitates with several DANs and MBONs implicated in memory updating and attractive behaviour. Finally we demonstrate that PD2a1/b1 receives axo-axonic synapses from MBON-α′2, an MBON implicated in appetitive memory retrieval.

We speculate that if PD2a1/b1 neurons are stimulated by appetitive odors (via PNs projecting to the LH) and converge with MBONs and DANs that mediate attractive behavior, PD2a1/b1 neurons may themselves play a role in olfactory-driven approach (Figure 7A-B). Therefore their decreased response to the CS+ after training may result in decreased attraction to the trained odor (Figure 7B) resulting in net avoidance of the CS+. To examine this, we attempted to stimulate PD2a1/b1 optogenetically in a place preference assay (Aso et al., 2014b) but due to off-target expression in the GAL4 and split-GAL4 lines we failed to observe a consistent phenotype (data not shown). To test this model, future studies will need to refine genetic control of PD2a1/b1 and clarify the connectivity between PD2a1/b1 and MBONs/DANs.

What function could this MB-to-LH connectivity fulfill? One possibility is to provide a means for learned and innate behaviors to interact in conflicting situations. MB-to-LH connectivity may also constrain what flies can learn through hard-wired circuitry in the LH (Jefferis et al., 2007; Marin et al., 2002; Wong et al., 2002). One recent study has implicated the MB in the integration of context and internal state (Lewis et al., 2015) during CO_2_ olfactory processing, implying a memory-independent role for the MB in olfactory processing. In this study we also identify potential contacts between PD2a1/b1 and DANs and MBONs. Moreover, similar potential convergence has been identified previously for two other LH output neurons (Aso et al., 2014b), for which no functional attribution has been reported so far. These data and our new study provide evidence that the MB and LH are interconnected in multiple ways and that the simple functional division of innate and learned olfactory behavior in the fly higher brain is an oversimplification.

The olfactory systems of both flies and mammals share the same basic blueprint (Su et al., 2009; Wilson, 2008). In mice, the piriform cortex is required for learning and memory (Choi et al., 2011), responds sparsely to odors (Stettler and Axel, 2009) and samples from the whole olfactory bulb (Miyamichi et al., 2011; Sosulski et al., 2011), similar to the MB. In contrast the olfactory amygdala is necessary and sufficient to instruct innate olfactory behavior (Root et al., 2014) and receives stereotyped input from the olfactory bulb (Miyamichi et al., 2011; Sosulski et al., 2011), drawing a comparison to the LH. Intriguingly, there are uncharacterised connections between the piriform cortex and olfactory amygdala (Schwabe et al., 2004) and we speculate that these connections may play a role in memory retrieval in the mammalian brain.

## Author Contributions

M-J.D., P.-Y.P, T.P., and G.S.X.E.J conceived the project and designed all experiments. M-J.D., G.B.-G, P.-Y.P, A.W., and A.H. performed experiments and analyzed data.., R.J.V.R., P.S., and G.S.X.E.J. carried out EM tracing and data analysis. S.F., Y.A., D.B., and G.R. contributed novel reagents and tools. The manuscript was written by M-J.D., A.S.B., P.-Y.P, T.P., and G.S.X.E.J.

## Acknowledgements

This work was supported by MRC LMB Graduate Studentships, Boehringer Ingelheim Fonds PhD Fellowships (M-J.D. and A.S.B) and a Janelia Graduate Research Fellowship (M-J.D.); ERC Starting (211089) and Consolidator (649111) grants and core support from the MRC (MC-U105188491) (to G.S.X.E.J); the Agence National de la Recherche funding of the MemoNetworks project (to P.-Y.P and T.P.) and the Labex Memolife for PhD fellowship to G.B.-G.; the Howard Hughes Medical Institute (A.W. and G.R.); and a Wellcome Trust Collaborative Award (203261/Z/16/Z to G.S.X.E.J., D.B., G.M.R.). This work was also supported by the Janelia Visiting Scientist Program. The FlyLight Project Team performed brain dissections, histological preparations and confocal imaging for the PD2a1/b1 Split-GAL4 characterisation, Polarity and MCFO data. Heather Dionne cloned the R71D08-LexA construct and created the split-GAL4 hemidriver transgenic lines. We thank the Bloomington Stock Center (NIH P40OD018537) and Matthias Landgraf for fly lines. We thank Paavo Huoviala for curating the behavioral significances of different glomeruli. We thank Fiona Love, Adam Heath, Philipp Ranft, Amelia Edmondson-Stait, Kimberly Meechan and Mahmoud Elbahnasawi for contributing 8 % of annotated synapses on PNs in the LH and Nadiya Sharifi for 4% of the MBON-α′2 cable. Finally we thank Marta Zlatic, Glenn Turner and his group, and the Jefferis and Préat groups for many insightful comments on the manuscript.

## Online Methods

### 1. Molecular Biology

The pBP-R71D08 gateway entry construct was a kind gift from Heather Dionne. The insert was transferred to the pBPLexA::p65Uw destination vector (Addgene) via a Gateway LR recombination (Invitrogen).

The enhancers used to create split-GAL4 hemidrivers were created based on annotations for PD2a1/b1 in a GAL4 expression pattern database (Jenett et al., 2012) as previously described (Aso et al., 2014a). The enhancer hemidriver lines were created as previously described (Pfeiffer et al., 2010).

All transgenic fly lines were generated by either Bestgene Inc or Genetic Services, Inc.

### 2. Drosophila Husbandry and Genetics

Standard techniques were used for fly stock maintenance and construction.For imaging and immunohistochemistry flies were raised at 25°C on standard Drosophila food. For MultiColor FlpOut (MCFO) experiments (Nern et al., 2015), the MCFO stock (see below) was crossed to either R37G11-GAL4, LH989 or LH991. Flies were collected after eclosion, transferred to a new food vial and incubated in a 37°C water bath for 20-25 minutes.

Transgenic lines used for behaviour (with the exception of optogenetic activation experiments) were outcrossed for five generations to a *w1118* strain in a wild-type Canton-Special (CS) background. For behavioral experiments (with the exception of optogenetic activation experiments) flies were raised at 18 °C and 60% humidity under a 12-hr:12-hr light-dark cycle.

For optogenetic activation experiments, flies were raised on standard Iberian containing yeast cornmeal and agar at 22 °C under a 12-hr:12-hr light-dark cycle. Behaviour was performed essentially as described (Aso et al., 2014b) For each vial of experimental flies, between 3-5 each of males and females were crossed on normal fly food supplemented with 1/500 all-trans-retinal (Sigma-Aldrich, MO, USA). Parental flies were allowed oviposit for 2-4 days before being tipped into a new vial, to control for population density of offspring. Upon eclosion females of the correct genotype were sorted on a cold plate and kept on fresh 1/250 all-trans-retinal food. Experiments were performed less than 3 days after cold sorting. All lines were grown at 22 °C in a covered box in a 12 hour light:dark cycle incubator. Optogenetic activation experiments were carried out at 50% humidity and 25°C.

All *Drosophila* strains used in this study are listed below:

- *w; +; 10XUAS-IVS-mCD8::GFP (attP2)*
- *w; LexAop2-Brp(d3)::mCherry (su(hw)attP5)* For labelling MBON-α2sc axons. A gift from Mathias Landgraf. (Christiansen et al., 2011)
- *w; Insite0089-GAL4* For labelling Av6a1 cell-type neurons. (Gohl et al., 2011)
- *w; +; R58G03-GAL4* (attP2) For labelling SIP cell-type neurons. (Jenett et al., 2012)
- *w; +; R37G11-GAL4 (attP2)* For labelling PD2a1/b1 cell-type neurons. (Jenett et al., 2012)
- *13xLexAop2-mCD8::GFP(su(Hw)attP8)* Reporter for the LexA system. Bloomington Stock Number: 32204 (Pfeiffer et al., 2010)
- *w; +; 20XUAS-IVS-mCD8::GFP (attP2)* Bloomington Stock Number: 32194
- *20xUAS-IVS-csChrimson::mVenus (attP18)* For optogenetic stimulation and split-GAL4 screening. Bloomington Stock Number: 55134 (Klapoetke et al., 2014)
- *w; LexAop-spGFP11* For GRASP experiments. Bloomington Stock Number: 58755 (Gordon and Scott, 2009)
- *w; +; UAS-spGFP1-10* For GRASP experiments. (Gordon and Scott, 2009)
- *w; R73B12-ADp65 (attP40); R37G11-DBD (attP2)* (this study). LH989, for labelling cell-type PD2a1/b1 neurons.
- *w; R29G05-ADp65 (attP40); R37G11-DBD (attP2)* (this study). LH991, for labelling cell-type PD2a1/b1 neurons.
- *w; +; R71D08-GAL4 (attP2)* For labelling the MBON-α2sc neurons. (Pfeiffer et al., 2010; • Séjourné et al., 2011)
- *w; +; UAS-Shi*^*ts1*^ For silencing neurons with a temperature shift (Bouzaiane et al., 2015; Kitamoto, 2001)
- *w; +; LexAop2-dTRPA1 (VK5)* For activating neurons with a temperature shift (Hamada et al., 2008; Pfeiffer et al., 2010)
- *w; +; LexAop2-TdTomato (VK5)* For confirming MBON-α2sc expression in landing site VK5 (Hamada et al., 2008; Pfeiffer et al., 2010)
- *w; R37G11-LexA (attp40)* For controlling PD2a1/b1 with the LexA system (Pfeiffer et al., 2010)
- *w; R71D08-LexA (attp40)* For controlling MBON-α2sc with the LexA system (Pfeiffer et al., 2010)
- *w;* +*; UAS-GCaMP3 (VK5)* For calcium imaging of odor responses after conditioning (Tian et al., 2009). Bloomington Stock Number: 32237
- *w, UAS-GCaMP6f (attP18)* For calcium imaging in thermal activation experiments (Chen et al., 2013)
- *w; pBPp65ADZpUw (attP40); pBPZpGAL4DBDUw (attP2)* Enhancerless split-GAL4 hemidrivers for behavioural control (Hampel et al., 2015; Pfeiffer et al., 2010)
- *w; +; 3xUAS-Syt::smGFP-HA (su(Hw)attP1), 5xUAS-IVS-myr::smGFP-FLAG (VK5)* Polarity and membrane marker (Aso et al., 2014a)
- *hsFlp2::PEST (attP3);+; 10XUAS-FRT>STOP>FRT-myr::smGFP-HA (VK00005), 10XUAS-FRT>STOP>FRT-myr::smGFP-V5-THS-10XUAS-FRT>STOP>FRT-myr::smGFP-FLAG (su(Hw)attP1)/ TM3, Sb* Reporter for MultiColor FlpOut (MCFO) (Nern et al., 2015)
- *w, LexAop2-CD8::GFP (su(Hw)attp8), 10xUAS-CD8::RFP (attp18)* For double labelling the membranes of R37G11-LexA and different DAN and MBON split-GAL4 lines. Bloomington Stock Number: 32229 (Liu et al., 2012)
- *w; R24E12-ADp65 (attP40); R52H01-DBD (attP2)* MB025B, split-GAL4 control of PAM-β’1 (Aso et al., 2014a)
- *w; R30G08-ADp65 (attP40); RTH-DBD (attP2)* MB032B, split-GAL4 control of PAM-β’2m (Aso et al., 2014a)
- *w; R20G03-ADp65 (attP40); R19F09-DBD (attP2)* MB018B, split-GAL4 control of MBON-α‘2 (Aso et al., 2014a)
- *w; R25D01-ADp65 (attP40); R19F09-DBD (attP2)* MB077B, split-GAL4 control of MBON-y2α‘1 (Aso et al., 2014a)
- *w; R12C11-ADp65 (attP40); R15B01-DBD (attP2)* MB002B, split-GAL4 control of MBON-β’2mp (Aso et al., 2014a)

**Table 1:** List of transgenic lines used for anatomical experiments. For behavioural experiments the genotypes are labelled in the figure.

| **<u>Figure Number</u>** | **<u>Genotype</u>** |
| --- | --- |
| Figure 1C | *w; LexAop2-Brp(d3)::mCherry (su(hw)attP5), R71D08-LexA (attP40)/+; R37G11-GAL4 (attP2)/10XUAS-IVS-mCD8::GFP (attP2)* |
| Figure 1F | *w; R73B12-ADp65 (attP40)/+; R37G11-DBD (attP2)/3xUAS-Syt::smGFP-HA (su(Hw)attP1), 5xUAS-IVS-myr::smGFP-FLAG (VK5)* |
| Figure 1G | *w, 20xUAS-IVS-csChrimson::mVenus (attP18)/w; +; R37G11-GAL4/+* |
| Figure 2A | *w, 20xUAS-IVS-csChrimson::mVenus (attP18)/w; R73B12-ADp65 (attP40)/+; R37G11-DBD (attP2)/+* |
| Figure 2B | *w, 20xUAS-IVS-csChrimson::mVenus (attP18)/w; R29G05-ADp65 (attP40); R37G11-DBD (attP2)* |
| Figure 5 | *w*; +; *UAS-GCaMP3 (VK5)/R37G11-GAL4 (attP2)* |
| Figure 6C | *w*, *LexAop2-CD8::GFP (su(Hw)attp8)* ,<br>*10xUAS-CD8::RFP (attp18)/w*;<br>*R24E12-ADp65 (attP40)/+*; *R52H01-DBD (attP2)/+* |
| Figure 6C' | *w*, *LexAop2-CD8::GFP (su(Hw)attp8)* ,<br>*10xUAS-CD8::RFP (attp18)/w*;<br>*R30G08-ADp65 (attP40)/+*; *RTH-DBD (attP2)+* |
| Figure 6C'' | *w*, *LexAop2-CD8::GFP (su(Hw)attp8)* ,<br>*10xUAS-CD8::RFP (attp18)/w*;<br>*R20G03-ADp65 (attP40)/+*; *R19F09-DBD (attP2)/+* |
| Figure 6E' | *w*, *LexAop2-CD8::GFP (su(Hw)attp8)* ,<br>*10xUAS-CD8::RFP (attp18)/w*;<br>*R12C11-ADp65 (attP40)/+*; *R15B01-DBD (attP2)/+* |
| Figure 6E'' | *w*, *LexAop2-CD8::GFP (su(Hw)attp8)* ,<br>*10xUAS-CD8::RFP (attp18)/w*;<br>*R25D01-ADp65 (attP40)/+*; *R19F09-DBD (attP2)/+* |
| Figure 6E''' | *w*, *LexAop2-CD8::GFP (su(Hw)attp8)* ,<br>*10xUAS-CD8::RFP (attp18)/w*;<br>*R25D01-ADp65 (attP40)/+*; *R19F09-DBD (attP2)/+* |
| Figure S1A | *w*, *13xLexAop2-mCD8::GFP(su(Hw)attP8)/w*;<br>*R71D08-LexA (attP40)/+* |
| Figure S1B | *w*; +; *R37G11-GAL4 (attP2)/10XUAS-IVS-mCD8::GFP (attP2)* |
| Figure S1C | *w*, *20xUAS-IVS-csChrimson::mVenus (attP18)/w*; *R73B12-ADp65 (attP40)/+*; *R37G11-DBD (attP2)/+* |
| Figure S2C | *w*; *LexAop2-Brp(d3)::mCherry (su(hw)attP5)*, *R71D08-LexA (attP40)/Insite0089-GAL4*; *+/10XUAS-IVS-mCD8::GFP (attP2)* |
| Figure S2D | *w*; *LexAop2-Brp(d3)::mCherry (su(hw)attP5)*, *R71D08-LexA (attP40)/+*; *R58G03-GAL4 (attP2)/10XUAS-IVS-mCD8::GFP (attP2)* |
| Figure S2E | *w*; *Insite0089-GAL4/+*; *10XUAS-IVS-mCD8::GFP/+* |
| Figure S6 | *w*; *R71D08-LexA<sub>p65</sub> (attP40)/+*; *LexAop2-TdTomato (VK5)/+* |

### 3. Immunohistochemistry

Throughout this study we used two different immunohistochemistry (IHC) protocols. Figures 1F, 2A-B, and S4 used Protocol 2 while all other IHC data was processed using Protocol 1.

#### Protocol 1

IHCs were performed as described (Ostrovsky et al., 2013). All specimens were mounted in Vectashield (H-1000) (Vector Laboratories, Burlingame, CA, USA).

#### Protocol 2

These IHCs were performed as described (Aso et al., 2014a). A full step-by-step protocol can be found at https://www.janelia.org/project-team/flylight/protocols. Following the IHC protocol the brains were fixed again in 4% Paraformaldehyde (Electron Microscopy Sciences, Hatfield, PA) for four hours at room temperature. The brains were mounted on poly-L-lysine-coated cover slips and dehydrated through a series of ethanol baths (30%, 50%, 75%, 95%, and 3 × 100%) each for 10 min. Following dehydration they were submerged in 100% Xylene three times for 5 minutes each. Samples were embedded in DPX (DPX; Electron Microscopy Sciences, Hatfield, PA).

**Table 2:** List of all antibodies used in this study, their catalog numbers and purpose.

| **<u>Antibody</u>**, **<u>Concentration</u>** | **<u>Catalog Number</u>** | **<u>Purpose</u>** |
| --- | --- | --- |
| Chicken anti-GFP, 1/1600 | Abcam, ab13970 | GFP or mVenus |
| Mouse anti-Brp, 1/20-1/30 | DSHB, University of Iowa, nc82 | neuropil |
| Mouse anti-GFP, 1/400 | Roche, 11814460001 | Reconstituted GFP (GRASP) |
| Rabbit anti-GABA, 1/200 | Sigma-Aldrich, A2052 | GABAergic neurons |
| Mouse anti-ChaT, 1/400 | DSHB, University of Iowa, 4B1 | Cholinergic neurons |
| Rabbit anti-DvGlut, 1/500 | Gift from Dion Dickman, University of Southern California, USA (Chen et al., 2017) | Glutamatergic neurons |
| Alexa-488 Goat anti-mouse IgG1, 1/1600 or 1/400 | Thermofisher, A-21121 | Secondary for GRASP (1/1600) and MCFO (1/400, V5) |
| Alexa-488 Goat anti-chicken, 1/1600 | Invitrogen, A-11039 | Secondary for GFP stain |
| Alexa-568 Goat anti-rabbit, 1/1600 | Invitrogen, A-21069 | Secondary for mCherry |
| Alexa-568 Goat anti-mouse, 1/1600 | Invitrogen, A-11004 | Secondary for nc82 |
| Alexa-633 Goat anti-mouse IgG1, 1/1600 | Invitrogen, A-21126 | Secondary for nc82 (FigS6) |
| Rat anti-FLAG tag, 1/200 | Novus Biologicals NBP1-06712SS | Primary for FLAG stain (MCFO and Polarity) |
| Rabbit anti-HA tag, 1/300 | Cell Signalling Technologies, C29F4 | Primary for HA stain (MCFO and Polarity) |
| Mouse anti-V5, 1/500 | AbD Serotec, DL550 | Primary for V5 (MCFO) |
| Cy5 Donkey anti-Rat, 1/150 | Jackson Immuno Research, 712-175-150 | Secondary for Polarity (FLAG) |
| Cy3 Goat anti-Rabbit, 1/1000 | Jackson Immuno Research, 111-165-144 | Secondary for Polarity and MCFO (HA) |
| Cy2 Goat anti-Mouse, 1/600 | Jackson Immuno Research, 111-165-144 | Secondary for Polarity (nc82) |
| Alexa-647 Donkey anti-Rat,<br>1/150 | Jackson Immuno Research,<br>712-605-153 | Secondary for MCFO (FLAG) |

### 4. IHC Image Acquisition

All images for IHC were acquired using a Zeiss 710 Confocal Microscope, essentially as described (Aso et al., 2014a; Cachero et al., 2010; Kohl et al., 2013). We used three modes of imaging: 20x, 40x and 63x.

For 20x imaging, whole mount brain and VNCs were imaged using a Plan-Apochromat 20x/0.8 M27 objective (voxel size = 0.56 × 0.56 × 1.0 μm; 1024 × 1024 pixels per image plane).

20x imaging was used for Figures 2A-B.

<u>For 40x imaging</u>, whole mount brains were imaged using an EC Plan-Neofluar 403/1.30 oil objective with 768 x 768 pixel resolution at each 1 μm, 0.6-0.7 zoom factor.

40x imaging was used for Figures 3A-C, S1A-B, S2E, S6.

<u>For 63x imaging</u>, whole mount brains were imaged using a Plan-Apochromat 63x/1.40 oil immersion objective (voxel size = 0.19 × 0.19 × 0.38 μm; 1024 × 1024 pixels). For certain images, tiles of regions of interest were stitched together into the final image (Yu and Peng, 2011).

63x imaging was used for Figures 1C, 1F-1G’’’, 5C-G, S1C, S4A.

### 5. Image Processing and analysis

To accurately label the presynapses of the LH-projecting MBONs, the 71D08-LexA driver was crossed to LexAop2-Brp(d3)::mCherry resulting in axon-specific labelling. For the region of the MBON under investigation (MBON-a2sc axons in the dorsal LH) a mask was created. Eight 71D08>Brp(d3)::mCherry brains were immunostained and registered onto a common template brain (JFRC2, http://www.virtualflybrain.org) using the nc82 counterstain. Image registration was carried out as described (Cachero et al., 2010) using the CMTK registration suite (https://www.nitrc.org/projects/cmtk). The boundary of the overlaid neurites for each region of interest from each brain was segmented manually as a mask in Fiji (https://fiji.sc/), using the Segmentation Editor function. The overlap was calculated against a large database of GAL4 expression patterns (Gohl et al., 2011; Jenett et al., 2012) also registered against JFRC2 (Manton et al., 2014) using the 

~~~
cmtk.statistics
~~~

 function in the open source nat (NeuroAnatomy Toolbox) package (https://github.com/jefferis/nat) for R (https://cran.r-project.org/). To control for the background signal of the brain we created a mask of the peduncle and performed the same overlap calculation for each GAL4 line. This peduncle overlap score was used normalise the MBON axon masks to produce the final overlap score for each GAL4 line. This allowed us to select lines with high signal-to-noise within the MBON masks and excluded expression patterns with Kenyon Cell expression which would confound behavioural analysis. The stacks GAL4 line expression patterns in the top 0.97 quartile were further analysed manually to identify Lateral Horn neurons.

For counting the number of cells in each line, images of R37G11-GAL4, LH989 and LH991 crossed to 20xUAS-csChrimson::mVenus (attp18) were used and cells manually counted.

MCFO brains were imaged in 63x mode (see above) and the stitched final image registered to the JFRC2013 template brain. Single neurons were manually annotated and segmented in 3D using Fluorender. For comparison with the data reconstructed from electron microscopy we automatically skeletonized MCFO image data using the filament editor tool provided by the image analysis software Amira 6.2.0, followed by manual editing. Morphological analysis was performed using NBLAST (see below). For analysis, skeletons were segregated into soma, primary neurite (the neurite that leads to the cell body), dendrite, primary dendrite (the neurite connecting the dendritic and axonal arbors) and axon by visual inspection using insight from our electron microscopy data. We isolated 5 cells from R37G11-GAL4, 13 cells from LH989 and 5 cells from LH991.All lines contained neurons which projected to the mushroom body calyx.

To examine overlap between PD2a1/b1 axons and MB neurons or PD2a1/b1 dendrites and PN axons, high resolution images (63x) of PD2a1/b1 split-GAL4 lines driving both a membrane and presynapse markers were segmented using Fluorender (http://www.sci.utah.edu/…are/127-fluorender.html). We compared this to published segmentations of the DANs and MBONs (Aso et al., 2014a). All data were registered to the JFRC2013 template brain (Aso et al., 2014a). For all cell-types in addition to the entire membrane stain, the axons and dendrites were segmented separately for at least n=2 well-registered brains. For each category of segmentation (dendrite-only, axon-only) we created a mask from their different samples by overlaying all the examples of each line. This was followed by contrast enhancement, gaussian blurring and auto-thresholding to create a mask. All image processing was performed using Fiji. Overlap comparisons for pairs of masks were compared in R using the cmtk.statistics function in the “nat” package.

Double labelling images performed with R37G11-LexA and different MBON and DAN Split-GAL4 lines were processed with a median filter using the despeckle command in Fiji. This was necessary to remove background due to the weak expression levels of the R37G11-LexA line.

### 6. Generation of split-GAL4 lines

Each split-GAL4 line consists of two hemidrivers, the p65ADZp in attP40 and the ZpGAL4DBD in attP2 (Pfeiffer et al., 2010). The lines were screened by combining these two hemidrivers with a copy of *20xUAS-IVS-csChrimson::mVenus (attP18).* The brains of females from each line were dissected and screened with an epifluorescence microscope. Split-GAL4 combinations with favourable expression patterns (sparse expression of PD2a1/b1) were double balanced to make a stable stock.

### 7. Behaviour: Olfactory Assays

For all behavior experiments, 0–2 day-old flies were transferred to fresh food vials the day before conditioning. Conditioning and tests of memory performance and of olfactory acuity were performed as described previously (Bouzaiane et al., 2015). Groups of 40-50 flies were trained with either one cycle of aversive training (single-cycle training), or five cycles spaced by 15 min inter-trial intervals (spaced training). During one cycle of training, flies were first exposed to an odorant (the CS+) for 1 min while 12 pulses of 5s-long 60V electric shocks were delivered; flies were then exposed 45 s later to a second odorant without shocks (the CS–) for 1 min. The odorants 3-octanol and 4-methylcyclohexanol, diluted in paraffin oil at 0.360mM and 0.325mM respectively, were alternately used as conditioned stimuli. The test of memory performance was performed in a T-maze. Flies were placed at the convergence point of two airflows interlaced with octanol or methylcyclohexanol from either arm of the T-maze. After 1 min in the dark, flies were collected from both arms of the T-maze for subsequent counting, yielding a score calculated as (N_CS+_ – N_CS-_)/ (N_CS+_ + N_CS-_). A single value of the performance index is the average of two scores obtained from two groups of genetically identical flies conditioned in two reciprocal experiments, using either odorant as CS+, and tested consecutively in the T-maze. Flies were maintained on food at all times, with the exception of during conditioning and memory test. Memory test occurred 10±5 minutes after conditioning, 3h±30 minutes after conditioning, and 24±1.5 h after conditioning to assay immediate memory, 3-h memory and long-term memory, respectively. For experiments involving neuronal blockade with Shi^ts^, the time courses of the temperature shifts are provided alongside each graph of memory performance, and periods of neurotransmission blockade are highlighted in red. Flies were transferred to the restrictive temperature (33°C) 30 min before the targeted time, to allow for acclimatization to the new temperature.

To measure innate odor avoidance towards 3-octanol or 4-methylcyclohexanol, naïve flies were placed at the convergence point of two airflows, one interlaced with 3-octanol or 4-methylcyclohexanol and the other from a bottle with paraffin oil only. The odor-interlaced side was alternated for successive groups that were tested. Odor concentrations used in this assay were the same as for memory assays. At these concentrations, both odorants are innately repulsive. The avoidance index was calculated the same way as the performance index in memory assays.

Memory scores are displayed as means ± SEM. A single value of the performance index is the average of two scores obtained from two groups of genetically identical flies conditioned in two reciprocal experiments, using either odorant as CS+, and tested consecutively in the T-maze. The indicated ‘n’ is the number of independent values of the performance index or avoidance index for each genotype. Memory graphs were subjected to statistical analysis using 1-way ANOVA followed by Newman-Keuls pairwise comparisons between the experimental group and its controls. ANOVA is robust against slight deviations from normal distributions or the inequality of variances if the sample sizes are similar between groups which was the case in our experiments. Therefore, we did not systematically test our data for normality or verify variance homogeneity prior to statistical tests, but we rather adopted a uniform analysis strategy for all our data ANOVA results are given as the value of the Fisher distribution F(x,y) obtained from the data, where x is the number of degrees of freedom between groups and y is the total number of degrees of freedom of the distribution. Statistical tests were performed using the GraphPad Prism 5 software. In the figures, asterisks illustrate the significance level of the t-test, or of the least significant pairwise comparison following an ANOVA, with the following nomenclature: *: p<0.05; **: p<0.01; ***: p<0.001; NS: not significant, p>0.05).

### 8. Calcium Imaging: Functional Connectivity

The genetically encoded GCaMP6f calcium reporter (Chen et al., 2013) (*UAS-GCaMP6f* in *attp18*) was driven by *R37G11 GAL4 (attP2)*. The thermosensitive cation channel dTrpA1 (Hamada et al., 2008) (*LexAop2-dTrpA1 VK00005*) was expressed in the V2 neurons by the 71D08-LexA driver (attP40). Female flies of the indicated genotypes were prepared for in vivo imaging as described above, and mounted on a custom-made chamber with controlled temperature through a Peltier cell and an analog electronic PID circuit. The baseline setpoint for the temperature was 20°C. Imaging was performed on the same setup as for olfactory responses, images were acquired at a rate of one image every 640 ms. During an acquisition with thermal activation, the setpoint of the temperature control circuit was shifted to 31°C for 30 s after a baseline recording of 10 s, and then back to 20°C. The measured risetime of the temperature from 20°C to 29°C was ∼8 s, and temperature reached 31°C within ∼11 s. For the temperature decrease was slower, taking ∼ 15s from 31°C to 22°C and ∼ 25s in total to decrease down to 20°C. For each fly, three such acquisitions were recorded, and the resulting time traces from visible hemispheres and from all these recordings were pooled and averaged. In R71D08LexA>LexAop2-TrpA1 flies, acquisitions with activation were alternated with acquisitions without activation as a permissive temperature control. The magnitude of activation was calculated as the mean of the time trace over a 20 s-time windows starting 5s after the change in temperature setpoint.

### 9. Calcium Imaging: Olfactory Responses

To monitor the olfactory responses in PD2a1/b1 neurons, the genetically encoded GCaMP3 calcium reporter (Tian et al., 2009) was driven by R37G11 GAL4 driver. We used a transgenic line carrying the UAS-IVS-GCaMP3-p10 construct inserted on the third chromosome in VK00005 (Bouzaiane et al., 2015; Tian et al., 2009). For *in-vivo* imaging, one female fly was prepared essentially as described previously (Séjourné et al., 2011). The fly was then placed under the objective lens (25x, 0.95 NA) of a confocal microscope under a constant airflow of 1.5 L·min-1. Images were acquired at a rate of one image every 128 ms. The emitted light was collected from transverse sections of the brain showing presynaptic terminals of PD2a1/b1 neurons. In general, both hemispheres could be recorded simultaneously. Olfactory stimuli were triggered by switching a valve to direct 30% of the total flow for 2 s through bottles containing odorants diluted in paraffin oil. Final dilution in the airflow was 1:2000. We recorded two series of responses to octanol and methylcyclohexanol, in alternating order, each separated by a 2 min interval, but only the first response to each odorant was kept for analysis. Data analysis was performed with Matlab software. For each recording, a ΔF/F0 time trace trace was calculated from an ROI surrounding the PD2a1/b1 projections. The baseline F0 value was calculated from the 2 s period preceding the switch of the valve. The response integral was calculated as the integral of the time trace during 10 consecutive timepoints following the onset of odor response (approx.. 2 s). The comparison of the response to a given odor between two groups, and of the response difference (Oct–Mch or Mch–Oct) between two groups, was performed using unpaired t-test.

### 10. Sparse Electron Microscopy Reconstruction and neuron identification

Neurons were reconstructed by ‘tracing’ in a full female adult Drosophila melanogaster brain volume (x,y,z resolution 4 nm x 4nm x 40 nm) that had been acquired by serial section transmission electron microscopy (Zheng et al., 2017), wherein the authors provide detailed sample preparation, EM acquisition and volume reconstruction protocols. Tracing aims to generate a neuronal skeleton that represents the branching of neurons and the locations of their synapses, rather than a volumetric reconstruction. Manual neuronal tracing through EM serial sections was performed in CATMAID (http://www.catmaid.org) (Saalfeld et al., 2009), a Web-based environment for working on large image datasets that has been optimised for tracing and online analysis of neuronal skeletons (Schneider-Mizell et al., 2016). Neuronal skeleton reconstruction was performed consistent with (Schneider-Mizell et al., 2016). Presynapses and postsynapses were annotated for all neurons traced in this study. Polyadic synapses, ubiquitous in *Drosophila* and common in insects (Meinertzhagen and O’Neil, 1991), were marked consistent with the criterion of other CATMAID-based *Drosophila* connectomic studies (e.g. (Zheng et al., 2017)). Briefly, synapses must have a clear presynaptic density, multiple vesicles in the vicinity of the density and a cleft between the pre- and postsynaptic membranes. Postsynapses were marked if they had a (though often unclear or faint) postsynaptic density or otherwise distinctive morphology in apposition to the synaptic cleft (Prokop and Meinertzhagen, 2006). Additionally, for PD2a1/1 neurons, the point at which microtubules ceased to be apparent in a branch was also annotated. Microtubules appear as thin dark filaments that flow contiguously from the cell body and terminate before the lowest order branches. Ambiguities and uncertainties in each neuron were flagged as it was traced, all neurons were subsequently and iteratively proofread and edited by an expert tracer until completion at least in the region of interest (see below). Gap junctions could not reliably be identified in this dataset.

MBONs-α2sc and MBON-α′2 were found by tracing downstream of extant reconstructed Kenyon cells (Zheng et al., 2017) within the appropriate mushroom body compartment (Aso et al., 2014a). Identity was verified with visual comparison to confocal stacks collected in (Aso et al., 2014a). Identification of PD2a1/b1 cell types began with tracing downstream of right-side MBON-α2sc. 23.95% of 1837 total outgoing connections from the right-side MBON-α2sc axon in the lateral horn were traced into 70 substantial neuronal arbors (> 300 μm of cable; data not shown). Visual inspection identified candidates for two PD2a1 neurons, which were traced to identification. This provided the location of the PD2 primary neurite tract (see Figure S7; nomenclature from Frechter et al. In Progress). In insect brains, the majority neuronal cell bodies are positioned outside of the neuropil proper, in the cortex, and invaginate the neuropil via a primary neurite before branching (Grueber et al., 2005). The primary neurite tract that a neuronal cell type takes is consistent between members of the type and between brains (SF and GSXEJ, in progress). No similar tract that might have also contained our neurons of interest could be found after thorough visual scanning through the EM data, nor was there any indication from NBLAST clustering of lateral horn neurons in the FlyCirciuit database (Chiang et al., 2011) or MCFO data that neurons similar to PD2a1/b1 could take multiple primary neurite tracts (data not shown). 185 neuronal profiles fasciculated within the PD2 tract, all of which were traced until their morphology made them an apparent PD2a1/b1 candidate or evidently not. Neurites for all candidates (34) were traced to or near ‘completion’ (see below). PD2a1 neurons must have 1) dendrite largely confined the the dorsal lateral horn, 2) primary neurite tract in the PD2 bundle, 3) an axon that circumvents around the mushroom body vertical lobe. Additionally PD2b1 neurons must have a process in the calyx. Two neurons met criterion 2 and 3, but were borderline on 1 and failed to receive similar projection neuron input to the 7 convincing members of the group, and were dropped from analysis. Identity was further verified by NBLAST (Costa et al., 2016) of reconstructed skeletons against MCFO data from this study and the FlyCircuit database (Chiang et al., 2011). Scores for our 7 putative PD2a1/b1 neurons were higher than for other candidate neurons in the PD2 tract and other MBON-α2sc targets (data not shown).

All 7 PD2a1/b1 neurons and the ipsilateral MBONs-α2sc and MBON-α′2 were fully traced ‘to completion’ *ex nihilo*, with synapse annotation. The contralateral MBONs-α2sc was traced to identification, but completed with the lateral horn. ‘Completion’ does not necessarily mean that absolutely all cable has been reconstructed and postsynapses and presynapses annotated, as a small minority of processes and connections may not have been resolved due to ambiguities in the image data. Many uniglomerular, excitatory projection neurons of the medial antennal lobe tract had been identified in the present EM volume, and traced outside the mushroom body calyx only to identification, not completion (Zheng et al., 2017). These PNs have since been reconstructed to completion in the LH (P.S, A.S.B et al., in preparation). For this study, we proofread, edited and annotated synapses for PN arbor in the right-side lateral horn for all 20 uniglomerular PN types in the vicinity of PD2a1/b1 dendrite and those determined to have significant overlap at a light level (data not shown). At first, one representative PN was chosen for each glomerulus that produced more than one uniglomerular, excitatory PN. If this PN was found to synapse onto PD2a1/b1 neurons, its sister cells were also completed within the lateral horn, as the morphology of sister PNs in the lateral horn are extremely similar (Jefferis et al., 2007).

### 11. Neuronal Skeleton Data Analysis

Neuronal skeleton data from CATMAID were analysed in R. Open source R packages for NBLAST (Costa et al., 2016), and R tools for accessing the CATMAID API are available on github by following links from jefferislab.org/resources. The catmaid and elmr R packages provide a bridge between a CATMAID server and the R statistical environment and bridging registration tools respectively. They include several add-on packages from the NeuroAnatomy Toolbox (nat see http://jeeris.github.io/nat/) suite enabling statistical analysis and geometric transformation of neuronal morphology. Further analysis relied on unreleased custom R code developed by A.S.B and G.S.X.E.J.

The elmr package provides tools for transforming data from the present EM whole female *Drosophila melanogaster* brain volume into different light level template brains for inspection of co-registered data. Neuronal skeleton reconstructions were brought from the EM brain space into the virtual flybrain template (http://www.virtualflybrain.org; dubbed JFRC2, the brain is divided into neuropils based on (Ito et al., 2014)) via the methods employed in Zheng and Lauritzen et al. 2017. FlyCircuit PD2a1/b1 neurons, identified through NBLAST clustering, were brought into the JFRC2 brain space using the Computational Morphometry Toolkit (https://www.nitrc.org/projects/cmtk/).

MBONs-α2sc, MBON-α′2 and PD2a1/b1 neurons were segregated into axon and dendrites using the sum of centrifugal and centripetal synapse flow centrality, algorithm (Schneider-Mizell et al., 2016), counting polyadic presynapses once. We verified that neurons were suitably polarized by calculating their axon-dendrite segregation index (Schneider-Mizell et al., 2016), which is a quantification for the degree of segregation of postsynapses and presynapses (0, totally unsegregated, 1, completely polarized). The mean±SD segregation index for PD2a1/b1 neurons was 0.27±0.09 indicating that these neurons are polarized but receive heavy axo-axonic modulation as well as outputting significantly in the lateral horn. MBON were highly polarized, for example right-side MBONs-α2sc had a segregation index of 0.72. Again we counted polyadic presynapses once, rather than using the number of outgoing connections these make, which would have been more expensive in terms of tracing time.

For morphological analysis of PD2a1/b1 neurons NBLAST (Costa et al., 2016) was performed on either the dendritic and/or the axonal arbors of neuronal skeletons. Primary neurite tracts and the primary dendrites connecting dendritic and axonal arbors were removed because their fasciculation, especially in the single EM brain space, made NBLAST less sensitive to dendritic and axonal differences. Clustering was performed using functions for hierarchical clustering in base R on euclidean distance matrices of NBLAST scores, employing Ward’s clustering criterion.

### 12. Data Presentation

All images of neuronal skeletons are shown in the JFRC2 brain space used by Virtual Fly Brain. Graphs were generated using the open source R package ggplot2 and related packages.

### 11. Data Availability

SWC files (Cannon et al., 1998) for the skeletonized multi-color flip-out data, and electron microscopy reconstructions for PD2a1/b1 neurons, and right-side MBON-α2sc and MBON-α′2 are available alongside the online version of this paper. Other data supporting the findings in this study are available upon request. A spreadsheet of glomeruli and published behavioural significance/functions are available upon request.

### 12. Code Availability

Core code described above is available by following links from jefferislab.org/resources. Further custom code can be made available upon request.

